# Improved TGIRT-seq methods for comprehensive transcriptome profiling with decreased adapter dimer formation and bias correction

**DOI:** 10.1101/474031

**Authors:** Hengyi Xu, Jun Yao, Douglas C. Wu, Alan M. Lambowitz

## Abstract

Thermostable group II intron reverse transcriptases (TGIRTs) with high fidelity and processivity have been used for a variety of RNA sequencing (RNA-seq) applications, including comprehensive profiling of whole-cell, exosomal, and human plasma RNAs; quantitative tRNA-seq based on the ability of TGIRT enzymes to give full-length reads of tRNAs and other structured small ncRNAs; high-throughput mapping of post-transcriptional modifications; and RNA structure mapping. Here, we improved TGIRT-seq methods for comprehensive transcriptome profiling by (i) rationally designing RNA-seq adapters that minimize adapter dimer formation, and (ii) developing biochemical and computational methods that remediate 5’- and 3’-end biases. These improvements, some of which may be applicable to other RNA-seq methods, increase the efficiency of TGIRT-seq library construction and improve coverage of very small RNAs, such as miRNAs. Our findings provide insight into the biochemical basis of 5’- and 3’-end biases in RNA-seq and suggest general approaches for remediating biases and decreasing adapter dimer formation.

## INTRODUCTION

High-throughput RNA sequencing (RNA-seq) has revolutionized biology and will become ever more powerful as new methods that address weaknesses and expand capabilities of current methods are developed (Mortazavi et al. 2008; Levin et al. 2010; Ozsolak and Milos 2011). A weakness of most current RNA-seq methods is their use of a retroviral reverse transcriptase (RT) to copy target RNAs into cDNAs, which are then be sequenced on different high-throughput DNA sequencing platforms (Head et al. 2014). Retroviral RTs have inherently low fidelity and processivity, and the extent to which these properties can be improved by protein engineering or in vitro evolution is limited by the retroviral RT structural framework (Hu and Hugh 2012).

To address this issue, we have been developing RNA-seq applications using the RTs encoded by mobile group II introns, bacterial retrotransposons that are evolutionary ancestors of introns and retroelements in eukaryotes (Mohr et al. 2013; Nottingham et al. 2016; Qin et al. 2016; Belfort and Lambowitz 2019). Unlike retroviral RTs, which evolved to help retroviruses evade host defenses by introducing and propagating mutational variations (Hu and Hughes 2012), group II intron RTs evolved to function in retrohoming, a retrotranposition mechanism that requires faithful synthesis of a full-length cDNA of a long, highly structured group II intron RNA (Lambowitz and Belfort 2015). Their beneficial properties for RNA-seq include high fidelity, processivity, and strand displacement activity, along with a proficient template-switching activity that is minimally dependent upon base pairing and enables the seamless attachment of RNA-seq adapters to target RNAs without RNA tailing or ligation (Mohr et al. 2013; Zhao et al. 2018). Thermostable group II intron RTs (TGIRTs) from bacterial thermophiles combine these beneficial properties with the ability to function at high temperatures (60-65°C), which help melt out stable RNA secondary structures that can impede reverse transcription (Mohr et al. 2013). A recent crystal structure of a full-length TGIRT enzyme (GsI-IIC RT, a form of which is sold commercially as TGIRT-III; InGex) in active conformation with bound substrates revealed that group II intron RTs are closely related to RNA-dependent RNA polymerases, as expected for an ancestral RT, and identified a series of novel structural features that may contribute to their high fidelity and processivity (Stamos et al. 2017). These features include more constrained binding pockets than retroviral RTs for the templating RNA base, 3’ end of the DNA primer, and the incoming dNTP, as well as a larger fingers region that enables more extensive contact with the template-primer substrate than is possible for retroviral RTs (Stamos et al. 2017).

GsI-IIC RT (TGIRT-III) has been used for a variety of applications, including comprehensive profiling of whole-cell, exosomal and plasma RNAs (Nottingham et al. 2016; Qin et al. 2016; Shurtleff et al. 2017; Boivin et al. 2018); quantitative tRNA-seq based on the ability of the TGIRT enzyme to obtain full-length end-to-end reads of tRNAs with or without demethylase treatment (Shen et al. 2015; Zheng et al. 2015; Qin et al. 2016); determination of tRNA aminoacylation levels (Evans et al. 2017); high-throughput mapping of post-transcriptional modifications by distinctive patterns of misincorporation (Katibah et al. 2014; Zheng et al. 2015; Shen et al. 2015; Shurtleff et al. 2017; Li et al. 2017; Safra et al. 2017); identification of protein-bound RNAs by RIP-Seq or CLIP (Katibah et al. 2014; Zarnegar et al. 2016); and RNA-structure mapping by DMS-MaPseq (Zubradt et al. 2017; Wang et al. 2018) or SHAPE (Mohr et al. 2018). A study comparing TGIRT-seq to benchmark TruSeq v3 datasets of rRNA depleted (ribodepleted) fragmented Universal Human Reference (UHR) RNA with External RNA Control Consortium (ERCC) spike-ins showed that TGIRT-seq: (i) better recapitulates the relative abundance of mRNAs and ERCC spike-ins; (ii) is more strand-specific; iii) gives more uniform 5’-to 3’-gene coverage and detects more splice junctions, particularly near the 5’ ends of genes, even from fragmented RNAs; and (iv) eliminates sequence biases due to random hexamer priming, which are inherent in TruSeq (Nottingham et al. 2016). Other recent studies have shown that TGIRT-seq more accurately depicts the quantitative relationship between mRNAs and structured small ncRNAs than other tested methods (Boivin et al. 2018) and eliminates artifacts due to RT mispriming in RNA-seq reactions (Shivram and Iyer 2018).

The TGIRT-seq method currently used for comprehensive transcriptome profiling (also referred to as TGIRT Total RNA-seq method) is outlined in Figure 1. This method uses the ability of TGIRT enzymes to template-switch directly from an artificial RNA template/DNA primer substrate containing an RNA-seq adapter sequence to the 3’ end of an RNA template, thereby coupling RNA-seq adapter addition to the initiation of cDNA synthesis (Mohr et al. 2013). For Illumina RNA-seq, the initial RNA template/DNA primer consists of a 34-nt RNA oligonucleotide containing an Illumina Read 2 sequence (R2 RNA) with a 3’ blocking group (C3 Spacer, 3SpC3) annealed to a 35-nt DNA primer (R2R DNA) that leaves a single nucleotide 3’ DNA overhang. The latter can base pair to the 3’ end of the target RNA, serving as a springboard for TGIRT template switching and initiation of cDNA synthesis (Mohr et al. 2013). To capture heterogeneous 3’ ends in a pool of RNAs, this single nucleotide 3’ overhang is an equimolar mixture of A, C, G, and T (denoted N) and is added in excess to the target RNA. After reverse transcription, a second RNA-seq adapter (R1R DNA; containing the reverse complement of an Illumina Read 1 sequence) is ligated to the opposite end of the cDNA by a single-stranded DNA ligation with thermostable 5’ App RNA/DNA ligase (New England Biolabs), and this is followed by minimal PCR amplification with primers that add Illumina capture sites and sequencing indices. By avoiding gel-purification steps, TGIRT-seq libraries can be generated rapidly (2 h up to the PCR step) from small amounts of starting material (1-2 ng input RNA).

**FIGURE 1.**
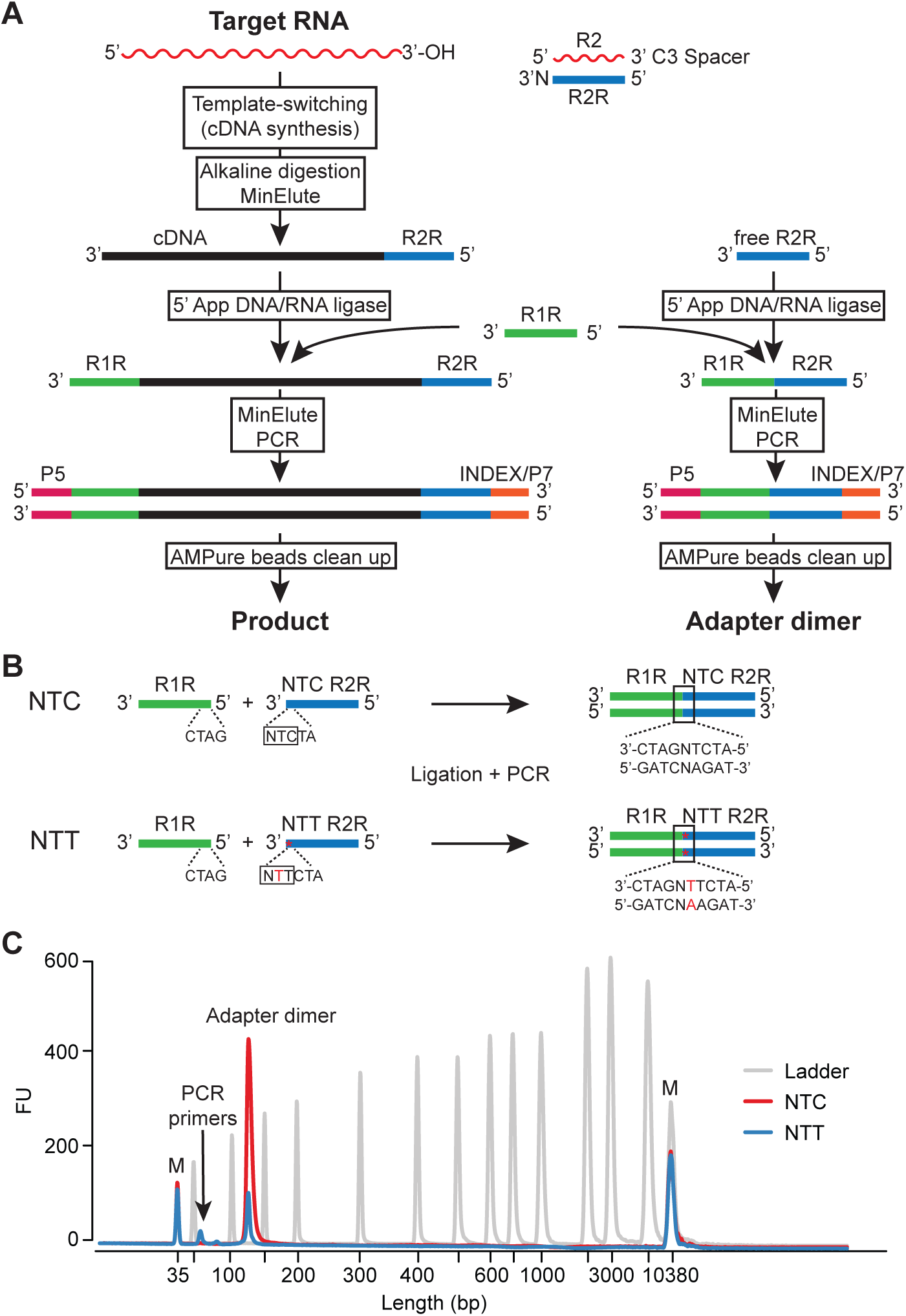
TGIRT-seq workflow and design of an improved R2R adapter that decreases adapter-dimer formation. (*A*) TGIRT-seq workflow. In the first step, TGIRT enzyme binds to an artificial template-primer substrate comprised of an RNA oligonucleotide containing an Illumina R2 sequence with a 3’-end blocking group (C3) annealed to a complementary DNA oligonucleotide (R2R) that leaves a single nucleotide 3’ overhang, which can direct template-switching by base pairing to the 3’ end of an RNA template. For the preparation of TGIRT-seq libraries from pools of RNAs, the DNA primer consists of a mixture of DNA oligonucleotides that leave A, C, G, and T 3’ overhangs (denoted N). After pre-incubation of the TGIRT enzyme with the target RNAs and template-primer (see Materials and Methods), template-switching and reverse transcription of an RNA template are initiated by adding dNTPs. The resulting cDNA with an R2R adapter attached to its 5’ end is incubated with NaOH to degrade the RNA template and neutralized with HCl, followed by two rounds of MinElute clean-up using the same column (Qiagen). A pre-adenylated oligonucleotide containing the reverse complement of an Illumina R1 sequence (R1R) is then ligated to the 3’ end of the cDNA by using thermostable 5’ App DNA/RNA ligase (New England Biolabs), followed by MinElute clean-up and 12 cycles of PCR amplification with primers that add indices and capture sites for Illumina sequencing. Unused R2R adapters that are carried over from previous steps are also ligated to the R1R adapter by using 5’ App DNA/RNA ligase (New England Biolabs), resulting in formation of adapter dimers (pathway at right), which are removed by AMPure beads clean-up prior to sequencing. (*B*) Taking into account known biases of the 5’ App DNA/RNA ligase (Jackson et al. 2014; Nottingham et al. 2016; Wu and Lambowitz, 2017), the R2R adapter used previously in TGIRT-seq (denoted NTC) was modified by inserting a single T-residue at position −3, creating a modified R2R adapter (denoted NTT), which decreases adapter-dimer formation. (*C*) Bioanalyzer traces comparing adapter-dimer formation using the previous NTC and improved NTT R2R adapters. 2 pmole of the each R2R adapter was ligated to 40 pmole of adenylated R1R adapter followed by 12 cycles of PCR according to the TGIRT-seq protocol and 1 round of clean-up with 1.4X AMPure beads to remove salt and PCR primers. The traces were obtained by using a 2100 Bioanalyzer (Agilent) with a high sensitivity DNA chip.

Unlike retroviral RTs, which have been studied extensively and optimized for biotechnological applications for decades, the recently introduced TGIRT enzymes and TGIRT-seq methods are potentially subject to considerable further improvement. In this regard, a weakness of the TGIRT Total RNA-seq method is the thermostable 5’ App RNA/DNA ligase used to attach the R1R adapter to the 3’ end of the cDNA, which introduces sampling biases for cDNA ends and produces unwanted adapter dimers by ligating the R1R adapter to residual R2R adapter carried over from previous steps. To avoid wasting reads, these adapter dimers are removed by AMPure beads clean-up of the library prior to sequencing (Fig. 1A), a step that can result in the differential loss of sequences corresponding to miRNAs and other very small RNAs, whose library products are close in size to adapter dimers (146 and 124 nt, respectively; Boiven et al. 2018). The problem is particularly acute with low abundance RNA samples where multiple rounds of AMPure beads clean-up may be required to sufficiently decrease the ratio of adapter dimers to small amounts of library products (Qin et al. 2016). Due in part to this limitation, miRNAs and other very small RNAs have been analyzed by an alternative TGIRT-seq method (the TGIRT CircLigase method), which was patterned after the method commonly used for ribosome profiling with retroviral RTs (Ingolia et al. 2009; Heyer et al. 2015). In the TGIRT-based version of this method, template-switching rather than RNA ligation is used to add a larger adapter containing both R1R and R2R sequences, and the resulting cDNAs with the linked R1R/R2R adapter are gel-purified and circularized with CircLigase for RNA-seq library construction (Mohr et al. 2013; Katibah et al. 2014).

Here, we used the previously determined ligation biases of the thermostable 5’ App RNA/DNA ligase (Jackson et al. 2014; Nottingham et al, 2016; Wu and Lambowitz 2017) to design an R2/R2R adapter with just a single nucleotide change that dramatically decreases the formation of adapter dimers, thereby improving the recovery of miRNA sequences and enabling the construction of TGIRT-seq libraries from smaller amounts of starting material. Additionally, using a miRNA reference set containing an equimolar mix of 962 human miRNAs, we systematically analyzed 5’- and 3’-end biases in TGIRT-seq introduced by the thermostable 5’ App RNA/DNA ligase and template switching, respectively, and developed biochemical and computational methods for ameliorating these biases. We found that the 5’-sequence biases introduced mainly by the ligase could be computationally corrected and that 3’-biases introduced by TGIRT template-switching could be corrected either computationally or by employing an altered ratio of 3’-overhang nucleotides in the R2 RNA/R2R DNA primer mix.

## RESULTS

### A single nucleotide change in the R2R adapter strongly decreases adapter dimer formation

Analysis of TGIRT-seq datasets obtained for fragmented UHRs or plasma DNA suggested that a major source of sequence bias is the DNA ligation step using the thermostable 5’ App DNA/RNA ligase, which has a preference for A or C and against U/T at position −3 from the 3’ end of the acceptor nucleic acid (Jackson et al. 2014; Nottingham et al. 2016; Wu and Lambowitz 2017). We noticed that the R2R adapter used currently for TGIRT-seq (denoted NTC based on its 3’ end sequence) has a C-residue at position −3 from its 3’ end, which favors the formation of R1R-R2R adapter dimers during the ligation step (Fig. 1B).

To address this difficulty, we designed a new R2R adapter (denoted NTT) in which a single T residue was inserted at position −3, thereby replacing the favored C at this position with a disfavored T, but leaving the remainder of the R2R sequence unchanged (Fig. 1B). This internal insertion requires a complementary insertion in the R2 RNA to maintain base pairing in the R2 RNA/R2R DNA heteroduplex. In a test reaction in which the NTC and NTT R2R DNAs were ligated to R1R DNA followed by PCR with primers that add Illumina indices and capture sites as in the TGIRT-seq protocol (Fig. 1A), this single nucleotide change decreased the recovery of the R1R-R2R adapter dimers by 82-89% (n = 3; Fig. 1C). The lower levels of adapter dimer formation enable the construction of TGIRT-seq libraries with fewer rounds of AMPure beads clean-up and better recovery of library products corresponding to miRNAs and other very small RNAs. These improvements in turn enable the construction of TGIRT-seq libraries from smaller amounts of starting material than is possible with the NTC adapter (0.05 pmole of a 40-nt RNA and 0.5 pmole of a 20-nt RNA with 96-98% and 88-99% lower levels of adapter dimers than the NTC adapter after 1 round of 1.4X AMPure beads clean-up; n = 3; Fig. 2).

**FIGURE 2.**
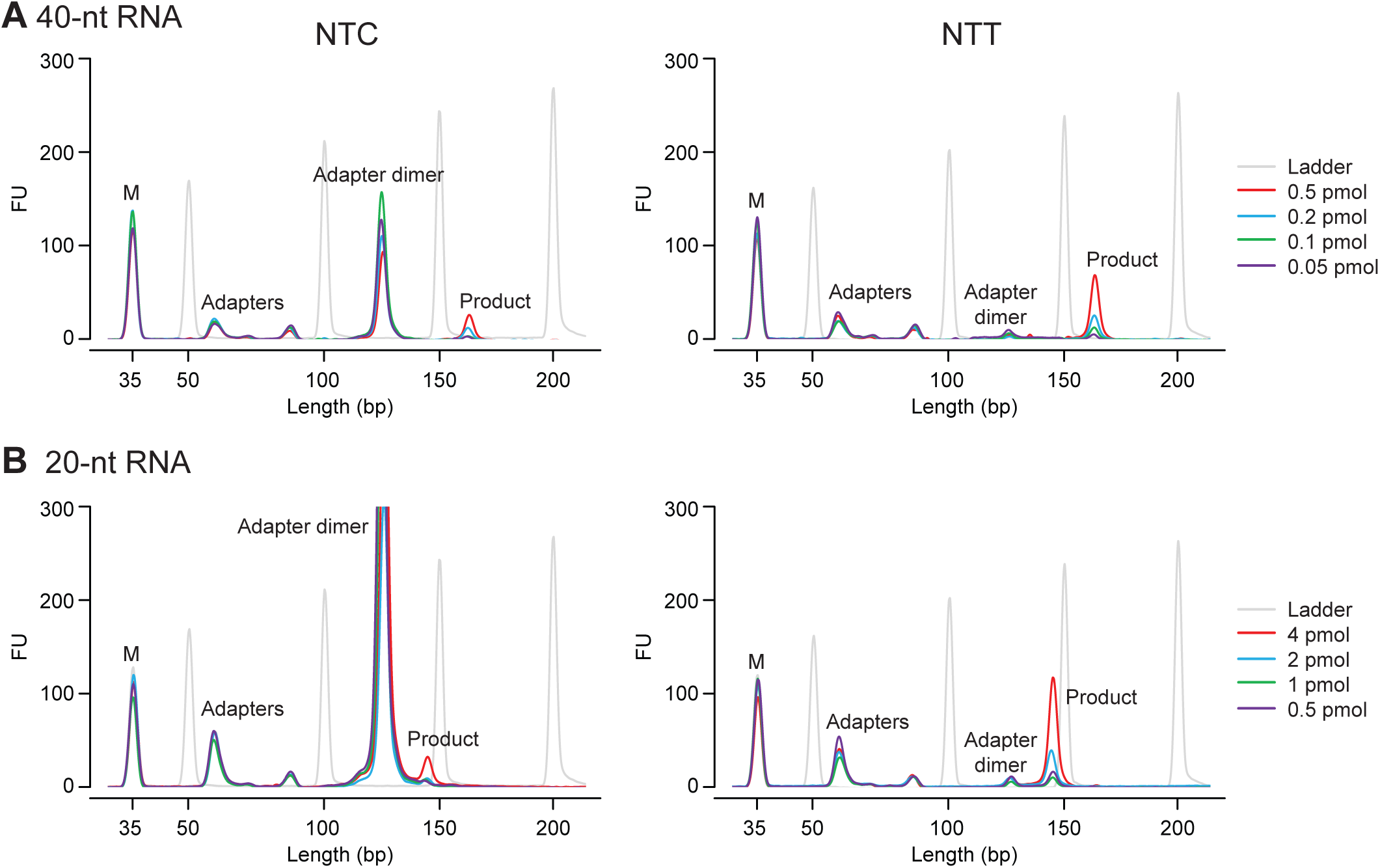
Bioanalyzer traces of TGIRT-seq libraries constructed from varying amounts of different-sized RNA oligonucleotides using either the NTC or NTT adapter. TGIRT-seq libraries were prepared from (*A*) 40-nt or (*B*) 20-nt RNA oligonucleotides using the workflow of Fig. 1A. After PCR for 12 cycles and one round of 1.4X AMPure beads clean-up, the libraries were analyzed on a 2100 Bioanalyzer (Agilent) using a high sensitivity DNA chip. M: internal marker.

### TGIRT-seq of ribodepleted, fragmented UHR RNA with ERCC spike-ins using the modified R2R adapter

To assess the performance of the NTT R2R adapter, we used it for TGIRT-seq of ribodepleted fragmented UHR RNA with ERCC spike-ins, as done previously for TGIRT-seq with the NTC adapter (Nottingham et al. 2016). The TGIRT-seq libraries were constructed in triplicate with one round of 1.4X AMPure beads clean-up to remove adapter dimers and sequenced on an Illumina NextSeq 500 to obtain 61-105 million 75-nt paired-end reads (Supplemental Table S1). The read-pairs were mapped to a human genome reference sequence (Ensembl GRCh38 modified to include additional rRNA repeats) by using an updated TGIRT-seq mapping pipeline (Materials and Methods). For comparison, raw sequencing reads from published TGIRT-seq datasets generated from similarly prepared fragmented UHR RNA samples using the NTC adapter (Nottingham et al 2016) were downloaded (NCBI SRA accession number SRP066009) and processed using the same bioinformatic pipeline.

The datasets obtained using the NTT adapter had mapping rates similar to those for the NTC adapter (84-86% and 84-89%, respectively), with similar proportions of the mapped reads mapping concordantly in the correct orientation to annotated genomic features (92-94%; Supplemental Table S1). Scatter plots comparing the representation of UHR RNAs in technical replicates obtained using the NTT and NTC adapters gave Spearman’s correlation coefficients (ρ) of 0.95-0.96 (Supplemental Fig. S1), and a histogram of the coefficients of variation of normalized counts from the replicates confirmed their similarly high reproducibility (94% and 92% of the protein-coding gene transcripts and spike-ins with normalized read count >10 have a coefficient of variation ≤25% for the NTT and NTC adapters, respectively, compared to 87% for TruSeq v3 in benchmark datasets (Li et al. 2014; Supplemental Fig. S2). Likewise, the normalized abundances (transcripts-per-million; TPM) of ERCC s-ins from the TGIRT-seq datasets correlated well with the expected spike-ins inputs (ρ = 0.98; Supplemental Fig. S3). The datasets obtained using the NTT and NTC adapters showed no substantial differences in the profiles of reads mapping to different genomic features (Fig. 3A,B), the distribution of reads between the sense and antisense strands of protein-coding genes (Fig. 3C), or the proportions of bases mapping to different regions of protein-coding genes (Fig. 3D).

**FIGURE 3.**
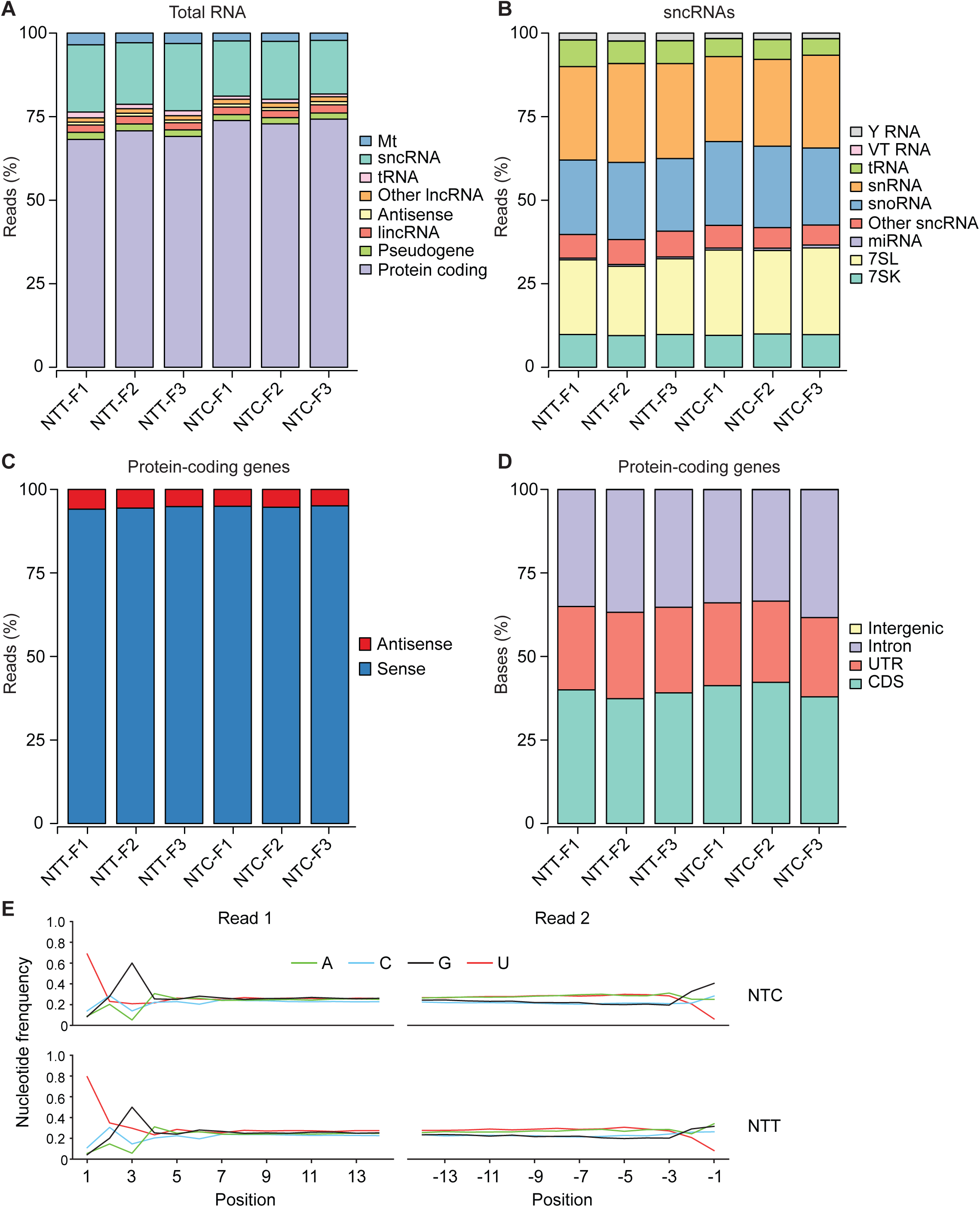
TGIRT-seq of ribodepleted fragmented UHR RNA with ERCC spike-ins using the NTT and NTC adapters. TGIRT-seq libraries were prepared in triplicate for each adapter and sequenced on an Illumina NextSeq 500 to obtain 58-105 million 75-nt paired-end reads, which were mapped to a human reference genomic (Ensembl GRCh38) modified to include additional rRNA repeats (Materials and Methods and Supplemental Table S1). The data were used to generate stacked bar graphs showing the percentages of: (*A*) read-pairs that mapped concordantly to the annotated orientation of different categories of genomic features; (*B*) small ncRNA reads that mapped to different classes of small ncRNAs; (*C)* protein-coding gene reads that mapped to the sense or antisense strand; (*D*) bases in protein-coding gene reads that mapped to coding sequences (CDS), introns, 5’- and 3’-untranslated regions (UTRs), and intergenic regions. The name of the dataset is indicated below. (*E)* Sequence biases at the 5’- and 3’-ends of RNA fragments in combined technical replicates of datasets obtained by TGIRT-seq of fragmented UHR RNAs with either the NTC or NTT adapters. Mapped reads from fragmented human reference RNAs using NTC (datasets NTC-F1 to -F3) or NTT (datasets NTT-F1 to F3) adapters were combined to calculate the nucleotides frequency at 14 positions at the 5’- and 3’-ends of the RNA fragments (positions +1 to +14 at the 5’ end of read 1 and −1 to −14 at the 5’ end of read 2, respectively).

To assess sequence biases in the TGIRT-seq libraries, we plotted aggregate nucleotide frequencies as a function of position from both the 5’- and 3’-ends of the sequenced RNA fragments (Fig. 3E). The plots show that 5’- and 3’-end sequences biases are similar for the NTT and NTC adapters, with the 5’ bias reflecting in large part the sequence preferences of the thermostable 5’ App DNA/RNA ligase (for C or A and against T at the −3 position of the cDNA, resulting in reciprocal biases for G or U and against A at position +3 from the 5’ end of the RNA), and the 3’ bias (for G and against U at position −1 from the 3’ end of the RNA) including a contribution from TGIRT-template switching. However, the contribution of the TGIRT-seq library preparation steps to the 5’- and 3’-end biases is difficult to assess from these datasets because the end biases could also include a contribution from the RNA fragmentation process (Parekh et al. 2016). Whatever the cause, nearly all of the sequence bias in libraries prepared from the fragmented human reference RNAs is confined to the first 3 positions from the 5’ and 3’ ends of the RNA fragments. Together, these results show that the NTT adapter performs similarly to the previous NTC adapter for analysis of ribodepleted fragmented whole-cell RNA preparations, but with fewer rounds of AMPure beads clean-up required to remove adapter dimers (1 round for the NTT adapter compared to 3 rounds in the previous libraries obtained with the NTC adapter; Supplemental Table S1).

### TGIRT-seq of a miRNA reference set and analysis of 5’- of 3’-end biases

To evaluate the performance of the NTT adapter in miRNA sequencing, we used it to construct TGIRT-seq libraries of a miRNA reference set containing an equimolar mixture of 962 human miRNA sequences (miRXplore Universal Reference; Miltenyi Biotech) and compared its performance to that of the NTC adapter tested in parallel. Libraries prepared using each adapter were constructed in triplicate, with the libraries constructed using the NTT adapter requiring 1 round of 1.4X AMPure beads clean-up prior to sequencing compared to 4 rounds for those constructed using the NTC adapter (Supplemental Table S2). The libraries were sequenced on an Illumina NextSeq 500 to obtain 10-16 million 2 x 75-nt paired-end reads, which were mapped to the 962 reference miRNA sequences. The proportion of uniquely mapped reads was higher for the NTT adapter than the NTC adapter (86-88% and 63-74%, respectively), possibly reflecting that the multiple rounds of AMPure bead clean-up required for the NTC adapter result in differential loss of *bona fide* miRNAs, which map uniquely, compared to larger aberrant products (*e.g*., resulting from multiple template switches), which do not map uniquely. Scatter plots comparing datasets for technical replicates gave ρ values of 1.00 for replicates with the same adapter and 0.94 for replicates with different adapters (Supplemental Fig. S4).

To assess sampling and end biases of the miRNAs in the TGIRT-seq datasets, we combined the 3 technical replicates for each adapter and compared the representation of miRNAs in the combined datasets to that in the reference set (Fig. 4A-C). Plots of the empirical cumulative distribution function (ECDF) of log_2_ normalized count for each miRNA in the reference set showed that the NTC and NTT adapters gave similar representations of different miRNA species (RMSE = 2.57 and 2.72, respectively; Fig. 4A-C, left panels). Additionally, the plots showed that, as from the fragmented UHR RNAs (Fig. 3E), much of the sequence bias in the TGIRT-seq datasets is confined to the first three nucleotides from the 5’- and 3’-ends of the miRNAs and is similar for the two adapters (Fig. 4A-C, middle and left panels).

**FIGURE 4.**
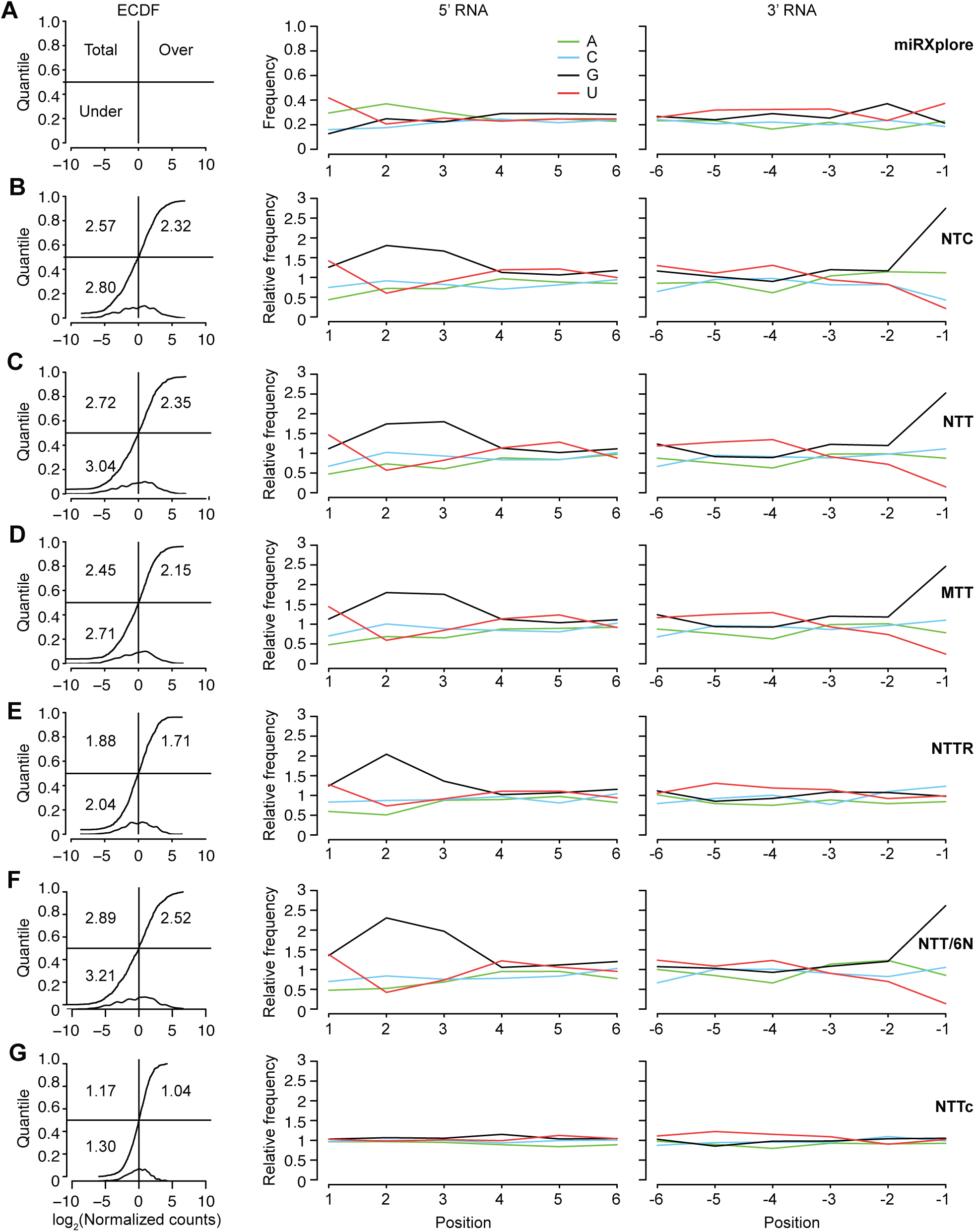
TGIRT-seq of the Miltenyi miRXplore miRNA reference set using the NTT or NTC adapters and different methods for mitigating 5’- and 3’-end biases. TGIRT-seq libraries were prepared from the Miltenyi miRXplore miRNA reference set containing 962 equimolar human miRNAs (Supplemental Table S2 and Materials and Methods). The datasets for each method (combined n = 3 for NTC, NTT, MTT, and NTTR; n =1 for NTT/6N) were used to plot both the empirical cumulative distribution (ECDF) function of the log_2_ median-normalized counts for each miRNA ranked from least to most abundant (left panels), and the abundance-adjusted nucleotide frequencies at the 5’-end (N+1 to N+6) and 3’-end (N-1 to N-6) for all miRNAs in the dataset relative to those in the miRNA reference set (middle and right panels). Only uniquely mapped reads were counted. The numbers within the ECDF plots for each method indicate the root-mean-square error (RMSE) for over-represented miRNAs (top right), under-represented miRNAs (bottom left), and all miRNAs (top left). (*A*) Miltenyi miRXplore reference set showing the ECDF plot layout (left panel) and 5’- and 3’-nucleotide frequencies for all miRNAs in the Miltenyi miRXplore reference set assuming equimolar concentrations of the 962 miRNAs. (*B*-*G*) ECDF plots (left panels) and plots of the ratio between the abundance-adjusted nucleotide frequencies at the 5’ and 3’ ends of miRNAs for TGIRT-seq and those in the miRNA reference set (middle and right panels) for datasets obtained using (B) the NTC adapter; (C) the NTT adapter; (D) a modified NTT adapter mix in which the 3’ A overhang is replaced with 3’ diaminopurine (MTT); (E) a modified NTT adapter mix with an altered ratio (6.6:0.4:1:1) of A, C, G, and T 3’ overhangs (NTTR); (F) the NTT adapter used together with an R1R adapter with six randomized nucleotides at its 3 end (NTT/6N); and (G) the NTT adapter after computational correction of 5’- and 3’-end biases (denoted NTTc).

Because the miRNAs in the reference set have known sequences, we could now more accurately assess the degree and cause of the bias introduced by TGIRT-seq than could be done with fragmented whole-cell RNAs. The 5’ bias includes but is not limited to the known sequence preferences of the 5’ App DNA/RNA ligase (*e.g.*, over-representation of G at position +3 of the RNA sequence compared to the reference set RNAs), while the 3’ bias due primarily to template-switching favors reference set miRNAs with a 3’ G residue and strongly disfavors miRNAs with a 3’ U residue (Fig. 4A-C, middle and right panels).

### Contribution of TGIRT-seq 5’- and 3’-end biases to miRNAs measurement errors

To quantify the contribution of TGIRT-seq 5’- and 3’-end biases to measurement errors for the miRNA reference set, we correlated the representation of each miRNA in the combined datasets obtained with the NTT adapter with its 5’- and 3’-end sequences. As the results for both the fragmented UHR RNAs (Fig. 3E) and the miRNA reference set (Fig. 4B,C, middle and right panels) showed that much of the bias is confined to the first 3 nucleotides from each end of the RNA, we focused on these positions. For this analysis, we defined over- and under-represented miRNAs as those whose log_10_ Counts-Per-Million (CPM) values were ≥1 standard deviation higher or lower than the mean log_10_ CPM for all of the miRNAs in the reference set (Supplemental Fig. S5). Principal component analysis (PCA) based on the first 3 nucleotides from the 5’- and 3’-end showed that the over- and under-represented miRNAs were almost linearly separable along the first principal component (PC1) of the PCA biplot (Fig. 5A).

**FIGURE 5.**
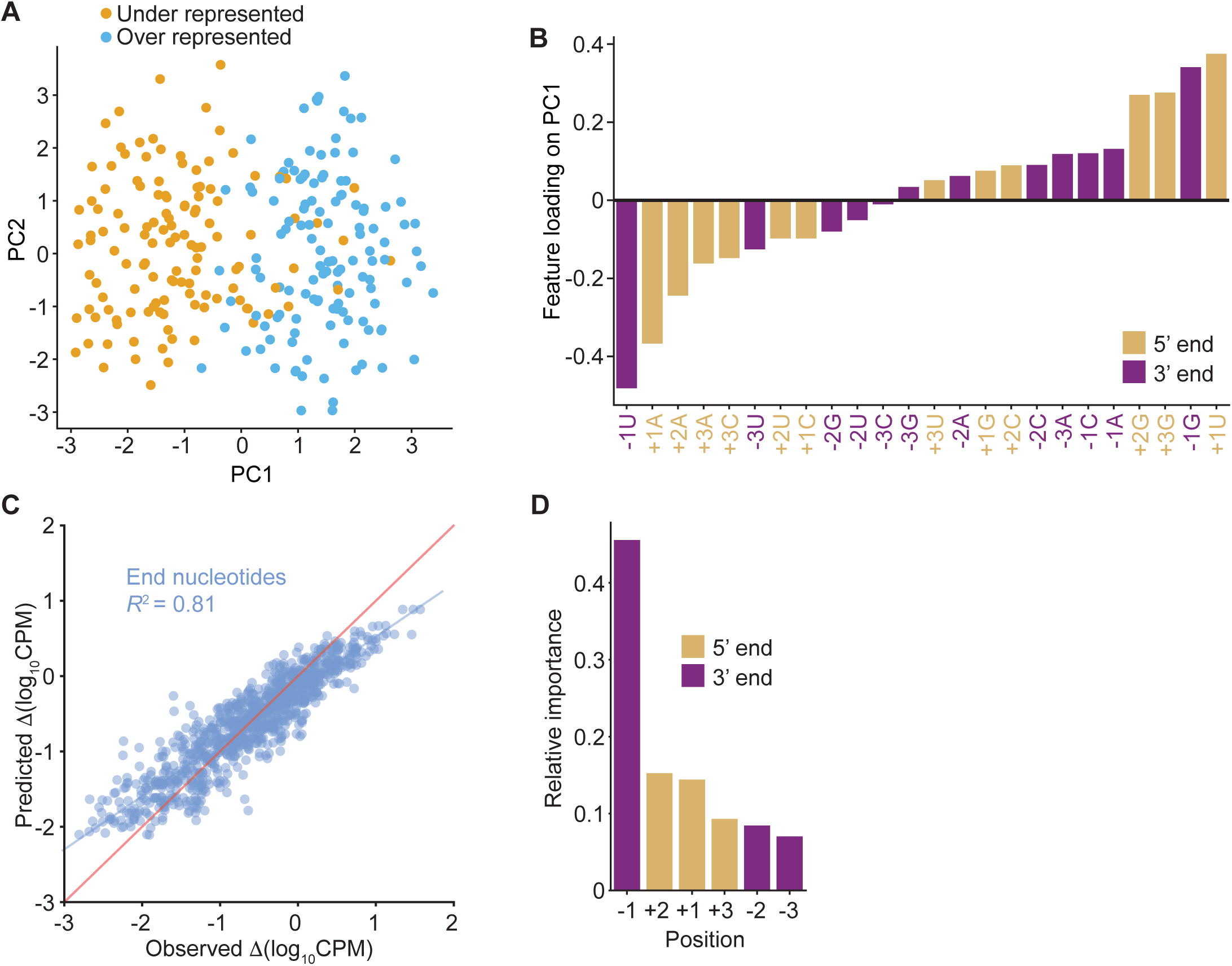
Effect of 5’- and 3’-end sequences on the representation of miRNAs in TGIRT-seq datasets. (*A*) Principal component analysis biplot for over- and under-represented miRNAs in TGIRT-seq of the Miltenyi miRXplore miRNA reference set in combined datasets for the three technical replicates obtained using the NTT adapter. The first three bases from the 5’- and 3’ ends of over- and under-represented miRNAs, defined as those whose log_2_CPM was at least one standard deviation greater or lower, respectively, than the mean log_2_CPM for all miRNAs in the reference set (Supplemental Fig. S5), were subject to principal component analysis. The first two principal components are shown. Each point indicates a miRNA, with over- and under-represented miRNAs colored as indicated in the Figure. (*B*) Relative importance of features of the first principal component. The fitted values from the first principal component are plotted for each base at each nucleotide position (feature) in ascending order. 5’- and 3’-end nucleotides are color coded as indicated in the Figure. (*C*) Random forest regression modeling of miRNA-seq quantification errors. A random forest regression model (*R*^2^ = 0.81) based on the first three 5’- and 3’-end positions was trained on the 962 miRNAs in the combined datasets for the 3 technical replicates obtained using the NTT adapter, and the predicted measurement errors (Δlog_10_CPM predicted by the model) were plotted against the observed measurement errors (Δlog_10_CPM obtained directly from sequencing data) for each miRNA. The blue line shows the fitted linear regression for the model, and the red line indicates hypothetical perfect prediction with slope = 1 and y-intercept = 0. (*D*) Relative importance of the position-specific preferences in TGIRT-seq. The relative importance of the 5’- and 3’-end positions from the random forest regression model were plotted in descending order. Each bar represents the relative importance of the indicated position color coded as indicated in the Figure.

To identify the contribution of different nucleotides to the miRNA recovery rate, we inspected the factor loadings on PC1 (Fig. 5B). This showed that 3 of the top 4 contributing factors in over-represented miRNAs were from 5’ positions with the most favored bases being 5’ +1U; +3G, and +2G (Fig. 5B, right side of plot). Moreover, 3 out of the top 4 contributing factors in under-represented miRNAs were also from the 5’ positions with the most disfavored bases being 5’ +1A, +2A, and +3A (Fig. 5B, left side of plot). However, the largest contributor for under-represented miRNAs and second largest for over-represented miRNAs was the 3’ terminal nucleotide (position −1), which favors a G residue and disfavors a U residue (Fig. 5B). By fitting the data to a random forest regression model, we found that the position-specific nucleotide preferences at the first three nucleotides from the 5’- and 3’-ends of the miRNA account for 81% (*R*^2^ = 0.81) of the measurement errors (Fig. 5C). A k-fold cross-validation test of the random forest regression model in which the 962 miRNAs were divided into 8 subgroups, each of which was tested with a model trained on the remaining subgroup, gave *R*^2^ values of 0.46 to 0.66 (Supplemental Fig. S6). By contrast, a model trained similarly using the internal positions +4 to +6 and −4 to −6 performed poorly (*R*^2^ = −0.06 to 0.06; Supplemental Fig. S6), confirming the importance of the first three 5’- and 3’-end positions.

As the random forest regression model predicts the continuous spectrum of miRNA measurement errors, we could use it to quantitatively assess the contributions of each of the first three 5’- and 3’-end position to the measurement errors (Fig. 5D). The results were generally consistent with the PCA, which identifies nucleotide combinations that separate over- and under-represented miRNAs. Thus, the −1 position was identified as having the greatest contribution to the bias, followed by the +2, +1 and +3 positions (Fig. 5D and Supplemental Fig. S6). A simple calculation summing the relative importance of the positions suggested that the 5’- and 3’-end biases contribute 40% and 60%, respectively, of the measurement errors due to end biases. However, we also found that nucleotides at some 5’- and 3’-end positions of the miRNAs in the reference set are correlated, in some cases with χ^2^-test −log_10_ p-values >10 (*e.g.*, 42% of the miRNAs with a disfavored A at +3 have a disfavored U at −1 position; Supplemental Fig. S7). This correlation raises the possibility that some of the apparent bias at 3’-end position −1 may reflect the 5’ adapter ligation bias rather than template-switching bias, consistent with this lower 3’-end bias seen in TGIRT-seq of a miRNA reference set using CircLigase instead of the 5’ App DNA/RNA ligase (Mohr et al. 2013, and see below).

### Biochemical and computational methods for remediating 5’- and 3’-biases in TGIRT-seq

Having investigated the sources of the 5’- and 3’-end bias in the TGIRT-seq protocol, we next explored biochemical and computational approaches for mitigating these biases. For the 3’ bias, we first thought that the preference for a G residue and against a U residue at position −1 might reflect the strength of the base-pairing interaction between that nucleotide and the 3’-overhang nucleotide of the DNA primer that is used to direct TGIRT-template switching, with a strong rG/dC base pair favored over a weak rU/dA pair. However, changing the 3’ A overhang in the NTT primer mix to a diaminopurine (denoted MTT) to enable a stronger base pair with 3 instead of 2 H-bonds to a 3’ U only slightly ameliorated this bias (RMSE decreased from 2.72 to 2.45; Fig. 4C,D).

We next tried an alternate approach based on previous findings that increasing or decreasing the proportion of a 3’-overhang nucleotide in the primer mix increases or decreases the recovery of miRNAs having a complementary 3’ end in the TGIRT-seq libraries (Mohr et al. 2013). We repeated this finding by constructing TGIRT-seq libraries from the miRNA reference set with a series of R2/R2R adapter mixes with higher proportions of 3’ A overhangs residues and lower proportions of 3’ C overhangs (Fig. 6) and found that we could almost completely eliminate the 3’ bias in TGIRT-seq of the miRNA reference set by using a primer mix with the ratio of 3’ overhang nucleotides A:C:G:T of 6.6:0.4:1:1 (denoted NTTR; RMSE = 1.88; Fig. 4E).

**FIGURE 6.**
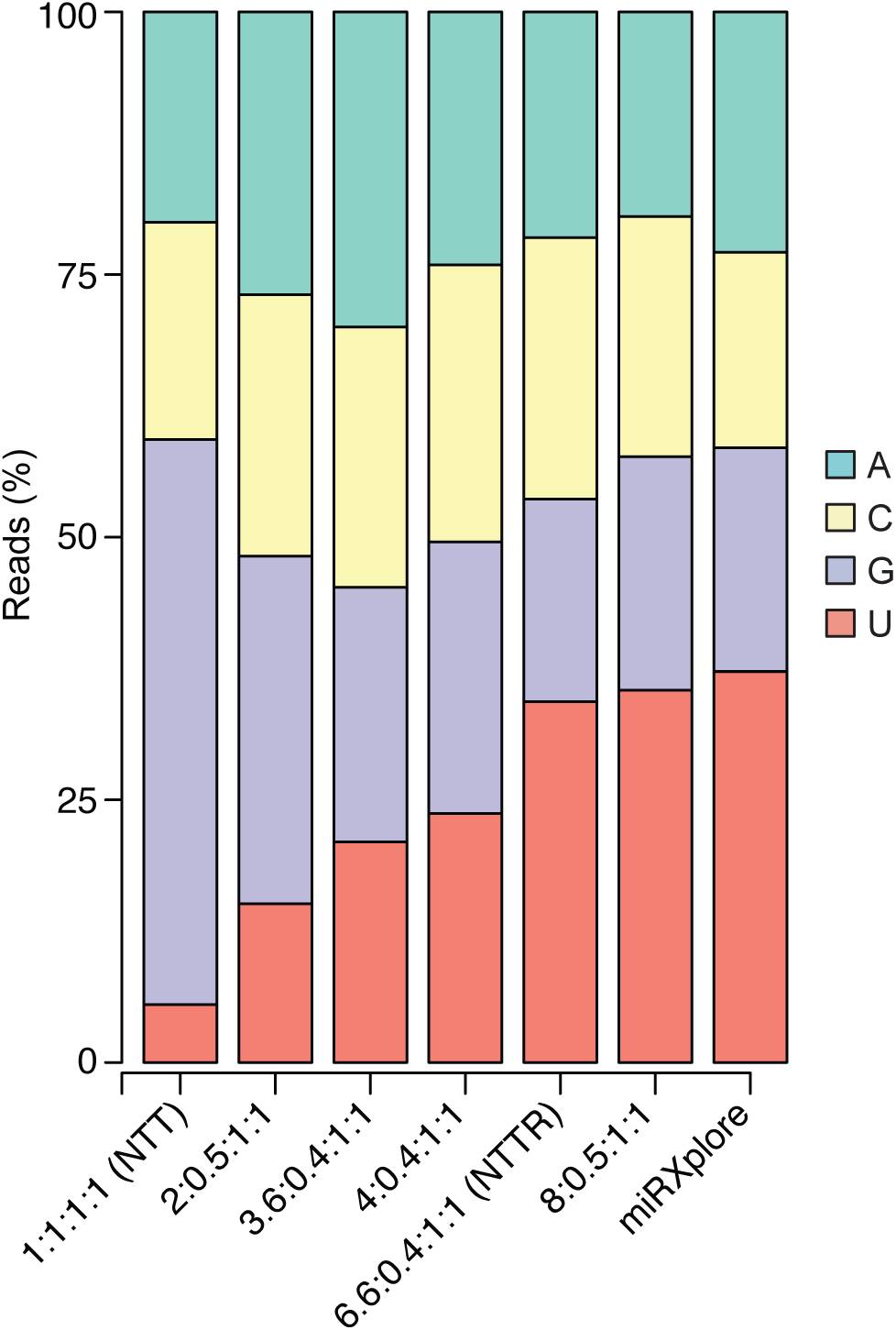
TGIRT-seq of the Miltenyi miRXplore miRNA reference set using R2 RNA/R2R DNA adapters with different ratios of the 3’-DNA overhang nucleotides. The stacked bar graphs show the percentages of miRNAs having A, C, G, and U 3’-end nucleotides, color coded as indicated in the Figure, in the datasets compared to those in the miRNA reference set (right). Only uniquely mapped reads were counted.

For the 5’-ligation bias, we noted that established small RNA-seq methods that employ T4 RNA ligases I and II to sequentially ligate DNA adapters to RNA 5’ and 3’ ends benefit from employing DNA adapters with four randomized nucleotides at the ligated ends (referred to as 4N protocols), with such adapters giving lower bias and better coverage at low sequencing depths than those with invariant sequences at their ends (Hafner et al. 2011; Jayaprakash et al, 2011; Zhuang et al. 2012; Giraldez et al. 2018). However, miRNA libraries prepared by TGIRT-seq with an R1R adapter containing six randomized 3’-end positions (denoted NTT/6N) did not decrease the ligation bias (RMSE = 2.89 compared to 2.72 for NTT with the R1R adapter without randomized nucleotides; Fig. 4F compared to Fig. 4C). This result may reflect that a major component of the ligation bias in methods that benefit from 4N adapters is thought to result from miRNA/adapter base-pairing interactions (referred to as “co-folding”), which can be ameliorated by providing more diverse adapter sequences (Hafner et al. 2011; Zhuang et al. 2012). By contrast, because TGIRT-seq employs a thermostable ligase for a single-stranded ligation of a DNA adapter to a cDNA at high temperature, the bias resulting from base-pairing interactions between the adapter and acceptor cDNA may already be minimal.

As an alternative for addressing the sampling biases in TGIRT-seq, we built a proof-of-concept bias corrector for TGIRT small RNA-seq using the random forest regression model described above (Fig. 5C,D) to correct the measurement errors due to 5’- and 3’-end biases. The bias corrector uses the first and last 3 nucleotides of each miRNA to predict the measurement errors, such that a corrected abundance can be computed by subtracting the predicted measurement error from the experimentally determined abundance for each miRNA. By employing this simple computational correction on the TGIRT-seq datasets obtained using the NTT adapter (denoted NTTc), both the 5’- and 3’-end biases were largely corrected, and the frequencies of miRNA 5’- and 3’-end nucleotides in the dataset closely approached those in the miRNAs in the reference set with the RMSE decreased to 1.17 (Fig. 4G).

### Comparison of TGIRT-seq to other miRNA sequencing methods

Fig. 7 compares TGIRT-seq of the miRNA reference set using the different methods of bias correction described above with published datasets obtained by using established small RNA library preparation methods on RNA samples containing the 962 miRNAs in the Miltenyi miRXplore reference set. Because some of the published datasets contain additional miRNAs, we created *in silico* subsamples containing only the 962 reference set miRNAs from each dataset for these comparisons. Fig. 7A shows saturation curves (*i.e.*, plots of the recovery of miRNAs with a read count of ≥10 as a function of sequencing depth), and Fig. 7B shows violin plots of the log_10_CPM values for the reference set miRNAs obtained by the different methods. The plots confirm the previous finding (Giraldez et al. 2018) that the 4N protocols perform better than other widely used small RNA-seq methods both in sampling miRNAs (reaching the plateau at smaller library sizes; Fig. 7A) and in obtaining expected log_10_CPM values (median closest to the red line) with smaller variance across the measured miRNA CPM values (shorter distance between the two ends of the tails; Fig. 7B). TGIRT-seq with NTTR adapter performed almost as well as the 4N protocols and better than TGIRT-seq with other adapters in the recovery of miRNA sequences as a function of read depth, likely reflecting that the altered ratio of R2R adapters 3’ overhangs improves the recovery or miRNAs with disfavored 3’ end sequences (Fig. 7A). Further, TGIRT-seq with the NTT or NTC adapters with computational correction (denoted NTTc and NTCc, respectively) performed slightly better than the 4N protocols in overall sampling bias and variance and substantially better than other commercial small RNA sequencing methods, including NEXTflex, TruSeq, CleanTag, and NEBNext (Fig. 7B). Based on a previously published dataset (Mohr et al. 2013; SRA accession number SRR833775), the TGIRT CircLigase method, employing TGRT-template-switching by TeI4c RT instead of GsI-IIC RT and a cDNA gel-purification step prior to circularization, performed about as well as the 4N protocols both in miRNA recovery as a function of sequencing depth and in overall sample bias and variance (Fig. 7A,B).

**FIGURE 7.**
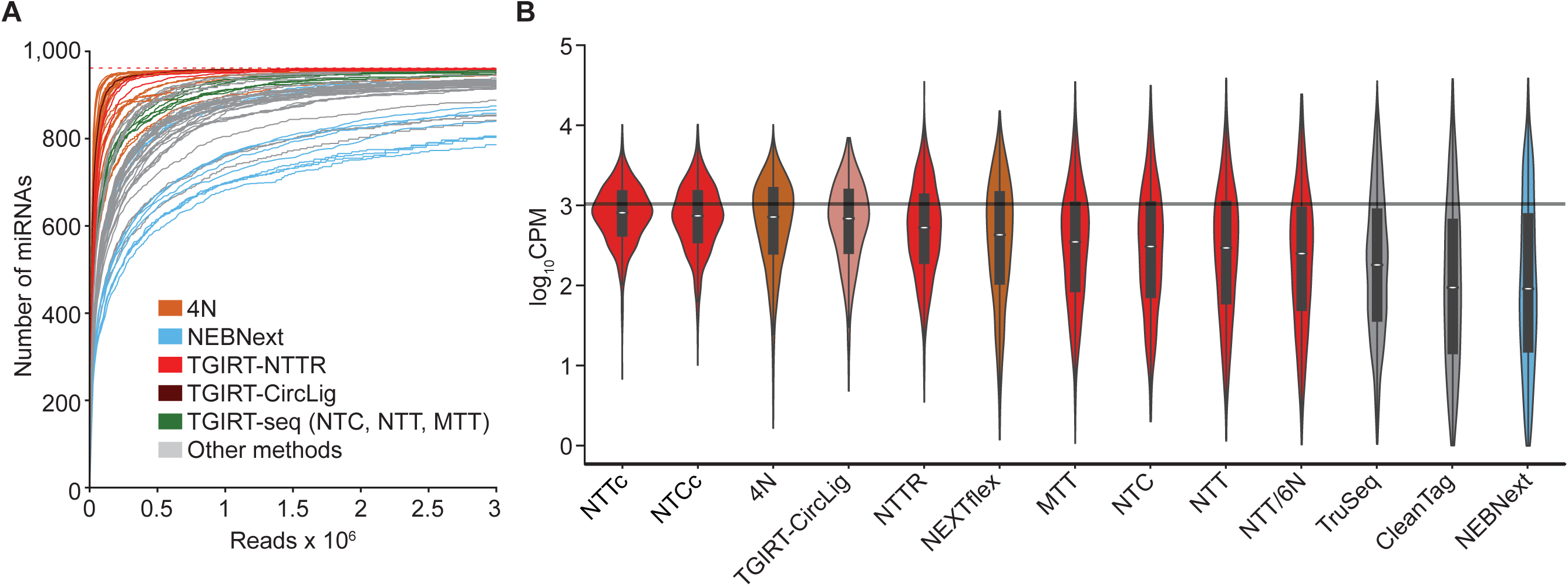
Saturation curves and differences in coverage for the 962 miRNAs in the Miltenyi miRXplore miRNA reference set for TGIRT-seq with or without different bias correction methods compared to published datasets for established small RNA-seq methods. For published datasets containing additional miRNAs, *in silico* subsamples containing only the 962 reference set miRNAs were used for the comparisons. (*A*) RNA-seq saturation curves. The curves show the number of reference set miRNAs with at least 10 reads at bins of 200 reads. As additional reads are included, the number of miRNAs with at least 10 reads increases. Curves are truncated at 3 million reads. The dotted red line at the top indicates the number of miRNAs in the Miltenyi miRXplore reference set. Each curve represents combined datasets, color-coded by the sequencing method as shown in the Figure for the best (4N ligation/NEXTflex; n = 24) and worst (NEBNext; n = 12) methods from the comparison of Giraldez et al. (2018), as well as TGIRT-seq (n = 3 for libraries prepared with the NTT, MTT, and NTC adapters), TGIRT-seq with the NTTR adapter (n = 3), TGIRT-seq with an R1R adapter containing six randomized 3’-end positions (NTT/6N; n=1), and the TGIRT-CircLigase method (n = 1; Mohr et al 2013). Other library preparation methods (gray lines) include NEBNext, TruSeq and CleanTag. (*B*) Violin plots of miRNA abundance in datasets obtained by different methods. The plots show the distribution of log_10_CPM for each miRNA in the reference set for each library preparation method (miRNA count = 2,886 for NTTc, 2,885 for NTCc, 23,088 for 4N ligation, 961 for TGIRT-CircLigase, 2,886 for NTTR, 5,522 for NEXTflex, 2,886 for MTT, 2,886 for NTC, 2,886 for NTT, 962 for NTT/6N, 30,757 for TruSeq, 3,815 for CleanTag, and 11,452 for NEBNext). NTTc and NTCc denoted TGIRT-seq datasets obtained using the NTT or NTC adapters that were computationally corrected using the random forest regression model trained with NTT dataset (Fig. 5C,D). The black horizontal line indicates the expected CPM values (CPM = 1,039.5) for each miRNA for a uniform distribution of 1,000,000 reads to 962 miRNAs (*i.e.*, unbiased sampling for each miRNA). The library preparation methods and correction methods are ordered from the lowest to highest deviation between the median CPM (white point within the violin) and the expected CPM. Median log_10_CPM values for each prep are connected by a blue line. The black boxes in the violins indicate the interval between first and third quartiles, with the vertical lines indicating the 95% confidence interval for each method.

### Factors other than end biases that may contribute to measurement errors in TGIRT-seq

To further investigate sources of bias in miRNA sequencing, we compared the over- and under-represented reference set miRNAs in datasets obtained by TGIRT-seq NTT and the 4N ligation protocols (Fig. 8). In agreement with the findings above, we found that most of the under- and over-represented miRNAs in TGIRT-seq compared to 4N are due to ligation and template-switching biases that could be substantially corrected computationally, so that the distribution of most of the reference miRNAs after correction largely matched that in 4N protocols (Fig. 8A,B). However, a small number of miRNAs remained substantially under-or over-represented in both TGIRT-seq and 4N protocols.

**FIGURE 8.**
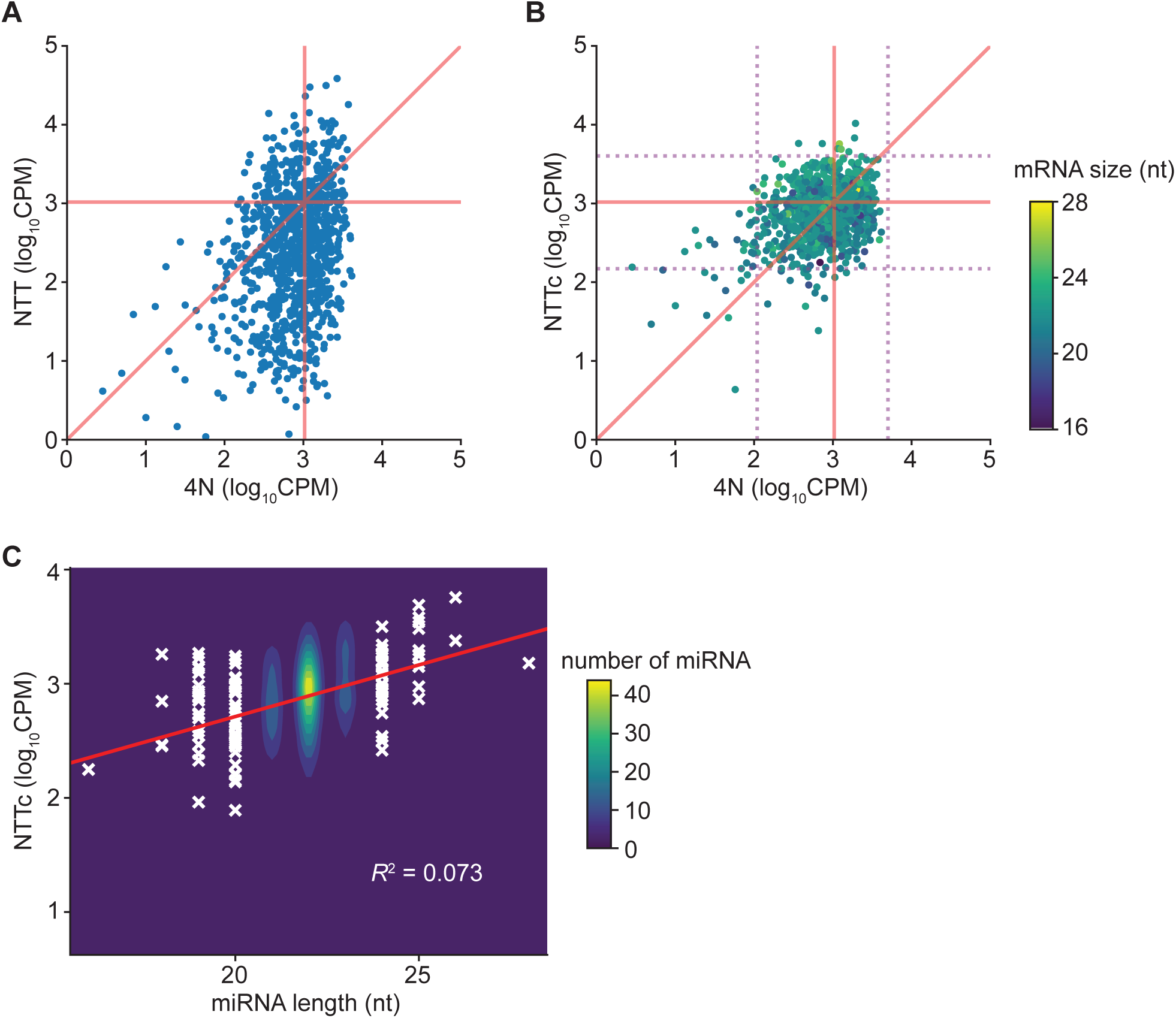
Representation of the Miltenyi miRXplore miRNA reference set in datasets obtained by TGIRT-seq with the NTT adapter before and after computational correction compared to representation of the same miRNAs in datasets obtained using 4N protocols. (*A*) miRNA representation for TGIRT-NTT versus 4N. Log_10_CPM values for each miRNA in combined TGIRT NTT datasets (n = 3) are plotted against those in combined datasets for 4N protocols (n = 24; Gilardez et al. 2018). Each point represents one miRNA. (*B*) The same comparison as (A) after computational correction of the TGIRT-seq NTT dataset using the random forest regression model (Fig. 5C,D). In (B), miRNAs are color-coded by their lengths (scale to the right). The purple dotted lines delineate 95% confidence intervals (2 standard deviations from the mean) of the miRNAs for 4N (vertical dotted lines) or NTT (horizontal dotted lines). The box delineates 892 miRNAs that lie within these confidence intervals and were used for comparison with over- and under-represented miRNAs in Fig. 9. The expected CPM values (CPM = 1,039.5 for each of the 962 equilmolar miRNAs) are indicated by horizontal and vertical orange lines for TGIRT-seq and the 4N protocols, respectively. The diagonal red line indicates cases where the CPM values from NTT are equal to those for 4N protocols. (*C*) Correlation between miRNA abundances and miRNA length. Two-dimensional kernel density estimation of the distribution for miRNA abundances and lengths (n = 962) is shown. The linear regression, with the equation: log_10_CPM= 0.09(miRNA size) + 0.9, is plotted as a red line, and miRNAs with length <21 or >23 nt are indicated as white crosses. The coefficient of determinant (*R*^2^) is indicated in the plot. The color scale indicates the numbers of miRNAs not shown as crosses.

To identify other factors that might contribute to the biased representation of these outlier miRNAs in TGIRT-seq, we defined over- and under-represented miRNAs as those whose log_10_CPM values after computational correction for end biases were ≥2 standard deviations higher (n = 8) or lower (n = 27) than the mean log_10_CPM, and then compared several potentially bias-inducing characteristics of these miRNAs to the remaining 927 more uniformly represented miRNAs (those in the center box in Fig. 8B). The compared characteristics included miRNA length, GC content, stability of potential secondary structure (self-fold free energy), potential of the miRNA cDNA with attached R2R adapter (resulting from the first step of TGIRT-seq) to base pair (co-fold) with the R1R adapter, and the numbers of unpaired (*i.e.*, free) 5’ and 3’ nucleotides in the most stable predicted secondary structure. Violin plots of the distribution of miRNAs in each group as a function of the compared characteristic showed that miRNA length was the only tested factor that contributes significantly to the under- or over-representation of these outlier miRNAs (Wilcoxon test p-values = 0.004 and 0.03, respectively; Fig. 9). However, for the larger group of 962 miRNAs, a plot of miRNA representation as a function of length showed only a weak correlation (*R^2^* = 0.073; Fig. 8C). The Violin plots confirmed that neither self-folding of the miRNAs nor co-folding of the miRNA cDNAs with the R1R adapter contributes significantly to the under-representation of the outlier miRNAs in TGIRT-seq (Fig. 9C,D).

**FIGURE 9.**
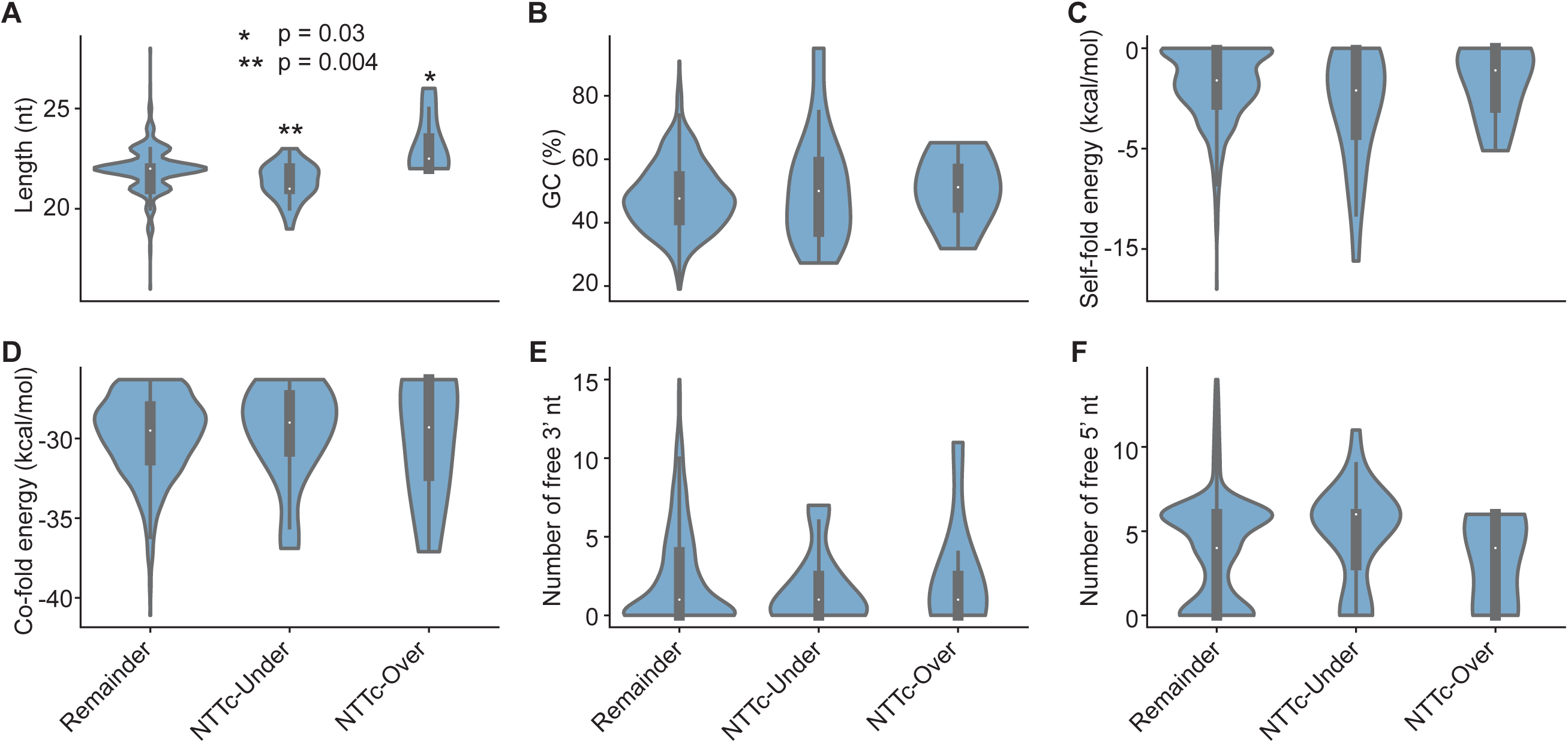
Factors other than end biases that may contribute to measurement errors in miRNA representation in TGIRT-seq. The figure shows violin plots comparing several potentially bias-inducing characteristics in over-represented (n = 8) or under-represented miRNAs (n = 27) defined as those with log_10_CPM values two or more standard deviations higher than the mean log_10_CPM compared to the remaining 927 miRNAs (those within the center box in Fig. 8B). The characteristics compared include: (A) miRNA length; (B) GC content; (*C*) the free energy of the most stable predicted secondary structure (self-fold energy) computed by the Vienna RNA package (Lorenz et al. 2011); (*D*) the predicted free energy of base pairing between the cDNA with attached R2R adapter (the product of the first step of TGIRT-seq) and the R1R adapter computed by Vienna RNA package (co-fold energy; Lorenz et al. 2011); (*E*) the number of unpaired (free) 3’ nucleotides in the predicted secondary structure; and (*F*) the number of unpaired (free) 5’ nucleotides in the predicted secondary structure. Asterisks on the top of the violins indicate significance of the difference between the outliers and remaining miRNAs determined by Wilcoxon test (*: p = 0.03; **: p = 0.004).

## DISCUSSION

By avoiding gel-purification steps, the TGIRT Total RNA-seq method enables the rapid construction of comprehensive RNA-seq libraries containing nearly all RNA biotypes from small amounts of starting materials with less overall bias than other transcriptome-profiling methods (Qin et al. 2016; Nottingham et al. 2016; Boivin et al. 2018). Here, we addressed two issues in TGIRT-seq library preparation, the disproportionate loss of miRNA sequences during AMPure beads clean-up of adapter dimers, and sampling biases resulting from 5’- and 3’-end sequences preferences in the ssDNA ligation and TGIRT template-switching steps.

First, to address the adapter dimer problem, we used the known sequence biases of the thermostable 5’ App DNA/RNA ligase employed for R1R adapter ligation to design an R2R adapter with a single nucleotide change that strongly decreases adapter dimer formation during TGIRT-seq library preparation (88-99% lower compared to the previous NTC adapter; Fig. 2). The redesigned R2R adapter (denoted NTT) decreases the number of rounds of AMPure beads clean-up required to remove adapter dimers, thereby increasing the recovery of very small RNAs and enabling the construction of TGIRT-seq libraries from smaller amounts of starting materials.

A previous approach for decreasing adapter dimer formation in RNA-seq protocols in which DNA adapters are ligated to both 5’- and 3’-RNA ends uses adapters with chemical modifications near the ligated ends of both adapters (Shore et al. 2016). These chemical modifications were hypothesized to inhibit ligation and impede subsequent reverse transcription when brought into close proximity in adapter dimers, but not when separated by a library insert. The authors carefully noted, however, that adapter dimer suppression was largely dependent upon the sequence of the adapters and that the same chemical modifications did not achieve the same degree of suppression with other adapter sequences (Shore et al. 2016). Our results extend these findings by showing that, at least for some ligases, small changes in adapter sequences based on analysis of ligase sequence preferences is by itself sufficient to strongly suppress adapter dimer formation without resorting to chemical modifications.

Next, we used TGIRT-seq of miRNA reference sets to analyze and correct 5’- and 3’-end biases. The 5’-end bias in TGIRT-seq is due in large part to sequence biases of the thermostable 5’ App DNA/RNA ligase used for single-stranded ligation of the R1R adapter to the 3’ end of the cDNA (Fig. 1A). We found that this bias could not be mitigated by using an R1R adapter with randomized nucleotides near its 5’ end, as in 4N ligation RNA-seq protocols, but could be corrected computationally by using a random forest regression model to give the same level of bias as in 4N protocols. The 3’-end bias in TGIRT-seq is confined largely to the 3’ nucleotide of the target miRNA, which base pairs with the 3’ overhang of the DNA primer mix during template switching. We first thought that this 3’ bias might reflect the relative strengths of the rG/dC and rU/dA base pair between the 3’ nucleotide of the miRNA and the 3’ overhang nucleotide of the primer. However, the 3’ bias could be only slightly ameliorated by substituting a diaminopurine 3’ overhang to enable a stronger base pair to a 3’ U, suggesting that it results largely results from nucleotide sequence preferences of the TGIRT enzyme. This 3’ bias could be almost completely remediated either by using primer mixes with a different ratio of 3’ A, C, G, and T overhang nucleotides to compensate for the sequence preferences of the TGIRT enzyme or computationally by using the random forest regression algorithm, which simultaneously corrects the 5’ bias (Fig. 4). The degree of computational correction that can be attained for TGIRT-seq is possible because sequences biases are almost entirely confined to the first 3 nucleotides from either end of the RNA template.

Surprisingly, although the computational correction for 5’ and 3’ end bias in TGIRT-seq and 4N ligation RNA-seq protocols address different factors, sequence bias in TGIRT-seq and adapter/miRNA co-folding in the 4N protocols (Giraldez et al. 2018), they achieve very similar degrees of overall correction in the datasets for miRNA reference sets, with relatively few outliers that are differentially corrected by one or the other method (Fig. 8). This likely reflects that the biases corrected by the two methods are orthogonal. The TGIRT-seq correction for 5’ end bias addresses sequencing preferences of the ligase, which are already minimal for the T4 RNA ligases used in the 4N protocols (Jayaprakash et al. 2011; Hafner et al. 2011; Zhuang et al. 2012; Fuchs et al. 2015; Giraldez et al. 2018), while the 4N correction addresses adapter/miRNA co-folding, which is likely already minimal in the high temperature ssDNA ligation in TGIRT-seq. Examination of outlier miRNAs after TGIRT-seq sequence bias correction indicates that miRNA length may be a contributing factor for under- and over-representation of some but not most miRNAs (Fig. 9). Although the recovery of miRNA sequences from pools of synthetic miRNAs, such as that used here, could in principal be used for bioinformatic approaches that fully correct all sources of biases, biological samples would likely behave differently from synthetic RNA pools tested at a single concentration *in vitro* (noted previously by (Giraldez et al. 2018)). Longer term, the 5’- and 3’-end biases in TGIRT-seq could be addressed further by using an alternative or modified ligase for 5’-adapter addition, while the 3’ bias and might be addressed by using a modified TGIRT enzyme. The recently determined crystal structure of full-length GsI-IIC RT in active conformation with bound substrates (Stamos et al. 2017) provides a platform for detailed analysis of the structural basis and possible alleviation of this 3’-end bias.

An attractive feature of the TGIRT Total RNA-seq method is that it enables the comprehensive analysis of different RNA size classes in a single RNA-seq experiment, enabling applications such as comparison of mRNA codon usage with cellular tRNA levels (Bazzini et al. 2016; Smith et al. 2018), co-expression of small ncRNAs and mRNAs encoding components of RNP complexes (Boiven et al. 2018), and analysis of tRNAs and tRNA fragments or mature, pre-, and pri-miRNA in the same RNA-seq (Nottingham et al. 2016; Qin et al. 2016; Burke et al. 2016; Wang et al. 2018). Previous work showed that the total RNA-seq protocol with TGIRT-III works well for quantitation of small RNAs down to ~60 nt (Boivin et al. 2018), and the introduction of the NTT adapter in the present work substantially improves the representation of miRNAs in the datasets. We note, however, that even with the NTT adapters, the recovery of miRNA sequences in the TGIRT Total RNA-seq method with GsI-IIC RTs (TGIRT-III) is less efficient than that of larger RNAs (Fig. 2), reflecting that miRNA library products may still be differentially lost at different clean-up steps in TGIRT-seq library construction (including the single round of Ampure beads clean-up that is still required to remove PCR primers) and that larger RNAs out compete very small RNAs (<60 nt) for reverse transcription by GsI-IIC RT in mixed-sized RNA preparations (Qin et al. 2016). For studies in which mature miRNAs are of primary interest, the latter issue could be minimized by introducing a size-selection step to obtain more uniformly sized RNA preparations and/or by employing an orthogonal approach, such as microarrays, RT-qPCR, or Firefly bead assay, to confirm inferences about miRNA abundance (Willenbrock et al. 2009; Chen et al. 2011; Wolter et al. 2014). Additionally, based on comparison of published datasets, we find that alternative TGIRT-CircLigase method, which includes a gel-purification step, performed similarly to 4N protocols in recovery of miRNA sequences as a function of sequencing depth as well as overall variance from the expected CPM values (Fig. 7), and at present, this may be the TGIRT method of choice for studies focused primarily on mature miRNAs. We also note that the another TGIRT enzyme, TeI4c RT, which has so far not been used extensively for RNA-seq, has significantly different properties than GsI-IIC RT, including the ability to synthesize even longer cDNAs and more uniform representation of RNAs <60 nt in mixed-sized RNA preparations (Qin et al. 2016). The numerous group II intron RTs identified by the sequencing of bacterial, archaeal, and organellar genomes may provide a rich resource for the identification of enzymes with even more beneficial properties for RNA-seq than those tested thus far.

## MATERIALS AND METHODS

### DNA and RNA oligonucleotides

The sequences of DNA and RNA oligonucleotides used in this work are summarized in Table 1. All oligonucleotides were purchased from Integrated DNA Technologies (IDT) in RNase-free HPLC-purified form. R2R oligonucleotides with different 3’ nucleotides were hand mixed prior to annealing to the R2 RNA oligonucleotide to obtain the desired ratio of single nucleotide 3’-overhangs (Nottingham et al. 2016; Qin et al. 2016). The NTT and NTC primer mixes contain an equimolar mix of R2R DNAs with 3’ A, C, G, and T residues. In the MTT primer mix, the R2R DNA with a 3’ A is replaced with a 3’ diaminopurine. In the NTTR primer mix, R2R DNAs with 3’ A, C, G, and T were mixed at a ratio of 6.6:0.4:1:1. Primer mixes with other ratios of 3’ nucleotides described in Results (Fig. 6) were prepared similarly.

**TABLE 1.**
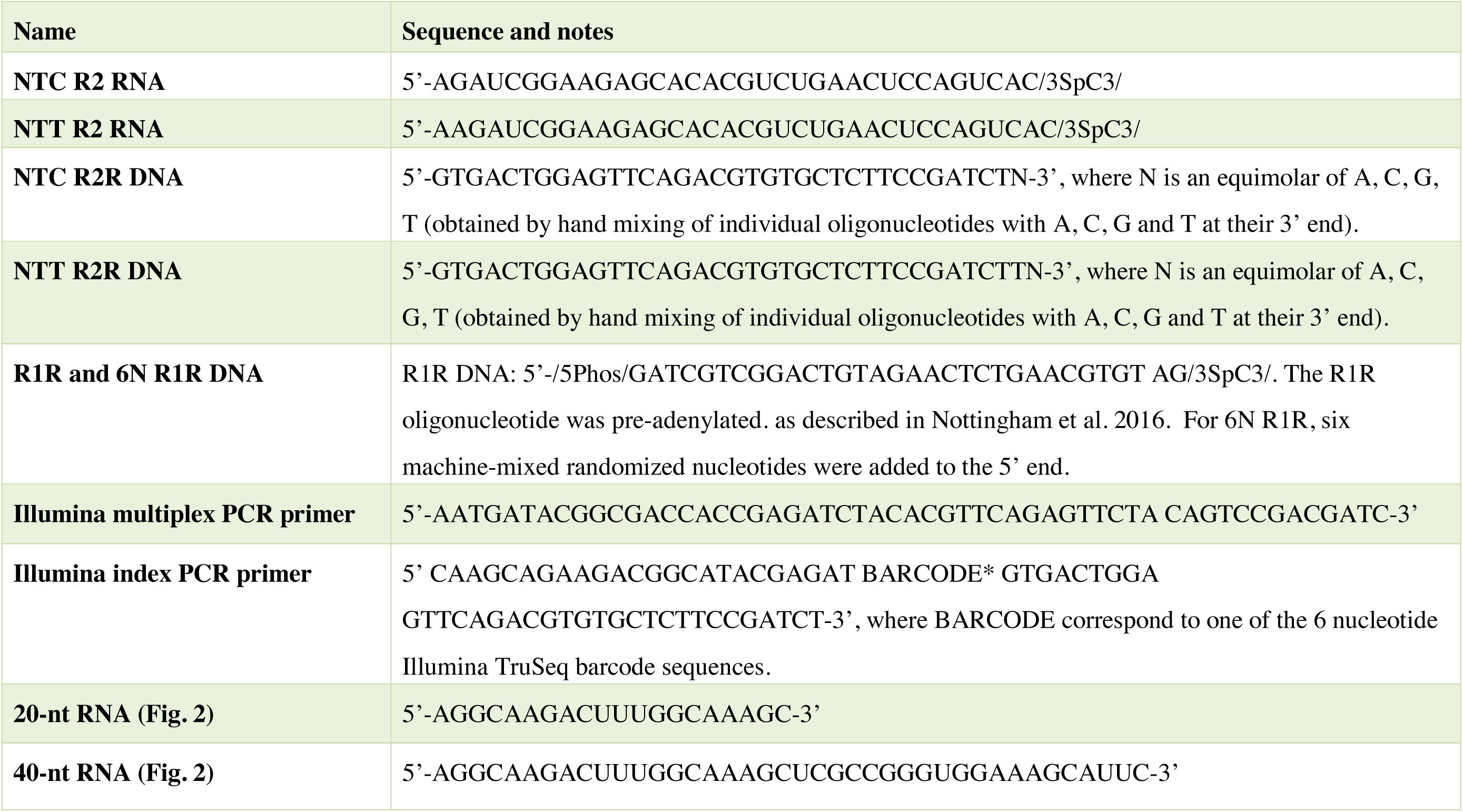
Oligonucleotides used in this study

### RNA preparations

The miRXplore miRNA reference set was purchased from Miltenyi Biotech. The RNA was dissolved in nuclease-free water (Invitrogen), adjusted to 1 μM, and aliquoted for storage at −80°C. Fragmented human reference RNA samples were prepared as described (Nottingham et al. 2016). 50 μl of Universal Human Reference RNA (UHR; Agilent) at 1 μg/μl was mixed with 1 μl of ERCC ExFold Mix 1 (Thermo Fisher Scientific; denoted ERCC spike-ins) prepared according to the provided protocol. 2 μl of the resulting UHR sample with ERCC spike-ins was ribo-depleted by using a Human/Mouse/Rat Ribo-zero rRNA removal kit (Illumina), fragmented to 70-100 nt by using an NEBNext Magnesium RNA Fragmentation Module (94°C for 7 min; New England Biolabs), and treated with T4 polynucleotide kinase (Epicentre) to remove 3’ phosphates that impede template switching (Nottingham et al. 2016). After each of the above steps, RNA was cleaned-up by using a Zymo RNA Clean & Concentrator kit, with 8 volumes of ethanol added to the input RNA to maximize the recovery of small RNAs (Nottingham et al. 2016). The fragment size range and RNA concentration were verified by using a 2100 Bioanalyzer (Agilent) with an Agilent 6000 RNA pico chip and aliquoted into 6 ng/3 μl portions for storage in −80°C.

### TGIRT-seq

TGIRT-seq libraries were prepared as described (Qin et al. 2016; Nottingham et al. 2016) using 6 ng of fragmented Universal Human Reference (UHR) RNAs with ERCC spike-ins or 50 nM Miltenyi miRXplore RNA prepared as described above. The template-switching and reverse transcription reactions were done with 1 μM TGIRT-III (InGex) and 100 nM pre-annealed R2 RNA/R2R DNA in 20 μl of reaction medium containing 450 mM NaCl, 5 mM MgCl_2_, 20 mM Tris-HCl, pH 7.5. The reactions were set up with all components except dNTPs, pre-incubated for 30 min at room temperature, a step that increases the efficiency of reverse transcription, and initiated by adding dNTPs (final concentrations 1 mM each of dATP, dCTP, dGTP, and dTTP). The template-switching reactions were incubated for 15 min at 60°C and then terminated by adding 1 μl 5 M NaOH to degrade RNA and heating at 95°C for 5 min followed by neutralization with 1 μl 5 M HCl and two rounds of MinElute column clean-up (Qiagen) to decrease the amount of unused R2R DNA adapter. The R1R DNA adapter was pre-adenylated by using an adenylation kit (New England Biolabs) and then ligated to the 3’ end of the cDNA by using thermostable 5’ App DNA/RNA Ligase (New England Biolabs) for 2 h at 65°C (Nottingham et al. 2016; Qin et al. 2016;). The ligated products were purified by using a MinElute Reaction Cleanup Kit and amplified by PCR with Phusion High-Fidelity DNA polymerase (Thermo Fisher Scientific; denaturation at 98°C for 5 sec followed by 12 cycles of 98°C 5 sec, 60°C 10 sec, 72°C 15 sec and then held at 4°C). The PCR products were cleaned up by using Agencourt AMPure XP beads (1.4X volume; Beckman Coulter), and sequenced on an Illumina NextSeq 500 instrument to obtain 2 x 75-nt paired-end reads.

### Bioinformatic analysis

Published TGIRT-seq data for identically prepared ribodepleted, fragmented UHR RNA plus ERCC spike-ins using NTC adapters were downloaded from NCBI (SRA accession number SRP066009). For UHR RNA plus ERCC spike-ins datasets, Illumina TruSeq adapters and PCR primer sequences were trimmed from the reads with cutadapt 1.16 (Martin 2011) (sequencing quality score cut-off at 20) and reads <15-nt after trimming were discarded. Reads were then mapped by using HISAT2 v2.0.2 with default settings to a human genome reference sequence (Ensembl GRCh38 Release 76) combined with additional contigs for 5S and 45S rRNA genes and the *E. coli* genome sequence (Genebank: NC_000913) (denoted Pass 1). The additional contigs for the 5S and 45S rRNA genes included the 2.2-kb 5S rRNA repeats from the 5S rRNA cluster on chromosome 1 (1q42, GeneBank: X12811) and the 43-kb 45S rRNA repeats that contained 5.8S, 18S and 28S rRNAs from clusters on chromosomes 13,14,15,21, and 22 (GeneBank: U13369). Unmapped reads from Pass 1 were re-mapped to Ensembl GRCh38 Release 76 by Bowtie 2 v2.2.6 (Langmead and Salzberg 2012) with local alignment to improve the mapping rate for reads containing post-transcriptionally added 5’ or 3’ nucleotides (*e.g.*, CCA and poly(U)), short untrimmed adapter sequences, or non-templated nucleotides added to the 3’ end of the cDNAs by the TGIRT enzyme (denoted Pass 2). The uniquely mapped reads from Passes 1 and 2 were combined using Samtools v1.8 (Li et al. 2009). To process multiply mapped reads, we collected up to 10 distinct alignments with the same mapping score and selected the alignment with the shortest distance between the two paired ends (*i.e.*, the shortest read span). In the case of ties between reads mapping to rRNA and non-rRNA sequences, the read was assigned to the rRNA sequence, and in other cases, the read was assigned randomly to one of the tied choices. Uniquely mapped reads and the filtered multiply mapped reads were combined and intersected with gene annotations (Ensembl GRCh38 Release 76) supplemented with RNY5 gene and its 10 pseudogene sequences, which were not annotated in this release, to generate the counts for individual features. Coverage of each feature was calculated by Bedtools (Quinlan and Hall 2010). To avoid mis-mapping reads with embedded sncRNAs, reads were first intersected with sncRNA annotations and the remaining reads were then intersected with the annotations for protein-coding genes, lincRNAs, antisense, and other lncRNAs. To further improve the mapping rate for tRNAs and rRNAs, we combined reads that were uniquely or multiply mapped to tRNAs or rRNAs in the initial alignments and re-mapped them to tRNA (Genomic tRNA Database and UCSC genome browser website) or rRNA (GeneBank: X12811 and U13369) reference sequences using Bowtie 2 local alignment. Because similar or identical tRNAs with the same anticodon may be multiply mapped to different tRNA loci by Bowtie 2, mapped tRNA reads were combined according to their anticodon (N = 48) prior to calculating the tRNA distributions.

For correlation analysis, RNA-seq datasets were normalized for the total number of mapped reads by using DESeq2 (Love and Huber 2014) and plotted in R. Reads that mapped to protein-coding genes were analyzed by Picard (http://broadinstitute.github.io/picard/) to calculate the percentage of bases in CDS, UTR, intron, and intergenic regions.

For datasets obtained for the Miltenyi miRXplore miRNA reference set, Illumina TruSeq adapters and PCR primer sequences were trimmed from the reads with cutadapt (sequencing quality score cut-off at 20) and reads <15-nt after trimming were discarded. Reads were then mapped with Bowtie2 using the local alignment with default settings to the Miltenyi miRXplore reference sequences. Uniquely mapped read with read lengths between 15 and 40 nt (86-88% of the mapped reads for the NTT adapter; Supplemental Table S2) were retrieved and used to calculate the counts table for 962 miRNAs. Counts from each dataset were median normalized, log_2_ transformed, and used to generate scatter plots, empirical cumulative distribution function (ECDF) plots, and nucleotide frequency plots in R. RMSE was calculated using log_2_ transformed median normalized counts.

### Correction of 5’- and 3’-end biases

miRNA sequence biases were analyzed with customized scripts using pysam (Li et al. 2009) and SciPy ecosystem (Jones et al. 2001). The deviations between the expected log_10_ miRNA abundance (log_10_CPM; 962 equilmolar miRNAs from Miltenyi miRXplore reference set) and measured log_10_ abundance were predicted from the first and last three bases for each miRNA using a random forest regression model implemented in R (Liaw and Wiener 2001) according to the following equation:

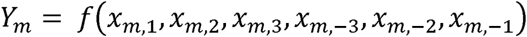

where *Y*_m_ indicates the difference between observed log_10_CPM and expected log_10_CPM for miRNA *m*. *f* indicates the random forest regression function, and *X*_m,i_ indicates the nucleotide of miRNA *m* at position *i*. Only the first 3 bases (*i* = 1 to 3) and the last 3 bases (*i* = −3 to −1) of each miRNA were considered. Correction of miRNA abundances was done by subtracting *Y*_m_ from the experimental log_10_CPM for each miRNA. All codes for miRNA modeling are deposited in GitHub at: https://github.com/wckdouglas/tgirt_smRNA.

### Comparison of TGIRT-seq of miRNAs to established small RNA-seq methods

miRNA count tables for 4N ligation, NEXTflex, TruSeq, NEBNext and CleanTag were downloaded from the National Center for Biotechnology Information (NCBI) Sequence Read Archive (SRA accession number SRP126845; Giraldez et al. 2018), and counts from the 962 Miltenyi miRXplore RNAs were extracted for the comparisons. Raw reads obtained by the TGIRT-CircLigase method (Mohr et al. 2013) were downloaded from NCBI (SRA accession number SRR833775), and aligned to the Miltenyi miRXplore reference sequences using Bowtie2 (settings local -D 20 -R 3 -N 0 -L 8 -i S,1,0.50 -k 5 --norc --no-mixed --no-discordant; Langmead et al. 2012) to generate a miRNA count table. miRNA counts from TGIRT-seq datasets and the downloaded datasets were normalized to CPM for the comparisons. The predicted RNA folding and co-folding patterns and energies were computed by the ViennaRNA package (Lorenz et al. 2011).

## DATA DEPOSITION

The TGIRT-seq datasets described in this manuscript have been deposited in the National Center for Biotechnology Information Sequence Read Archive under SRA accession number SRP168562.

## SUPPLEMENTAL MATERIAL

Supplemental material is available for this article.

## COMPETING INTEREST STATEMENT

Thermostable group II intron reverse transcriptases (TGIRT) enzymes and methods for their use are the subject of patents and patent applications that have been licensed by the University of Texas and East Tennessee State University to InGex, LLC. A.M.L. and the University of Texas are minority equity holders in InGex and some former Lambowitz laboratory members receive royalty payments from sales of TGIRT enzymes and licensing of intellectual property.

## ACKNOWLEDGMENTS

This work was supported by National Institutes of Health grant R01 GM37949 and Welch Foundation Grant F-1607. We thank our University of Texas at Austin colleagues Drs. Can Cenik and Vishwanath Iyer for comments on the manuscript. The authors acknowledge the Texas Advanced Computing Center (TACC) at the University of Texas at Austin for providing high performance computing resources that have contributed to the research results reported within this paper. URL: http://www.tacc.utexas.edu. The authors also acknowledge the Genome Sequencing and Analysis Facility (GSAF) and the Center for Biomedical Research Support (CBRS) at the University of Texas at Austin for providing sequencing services and computing resources.

---

**FIGURE S1.**
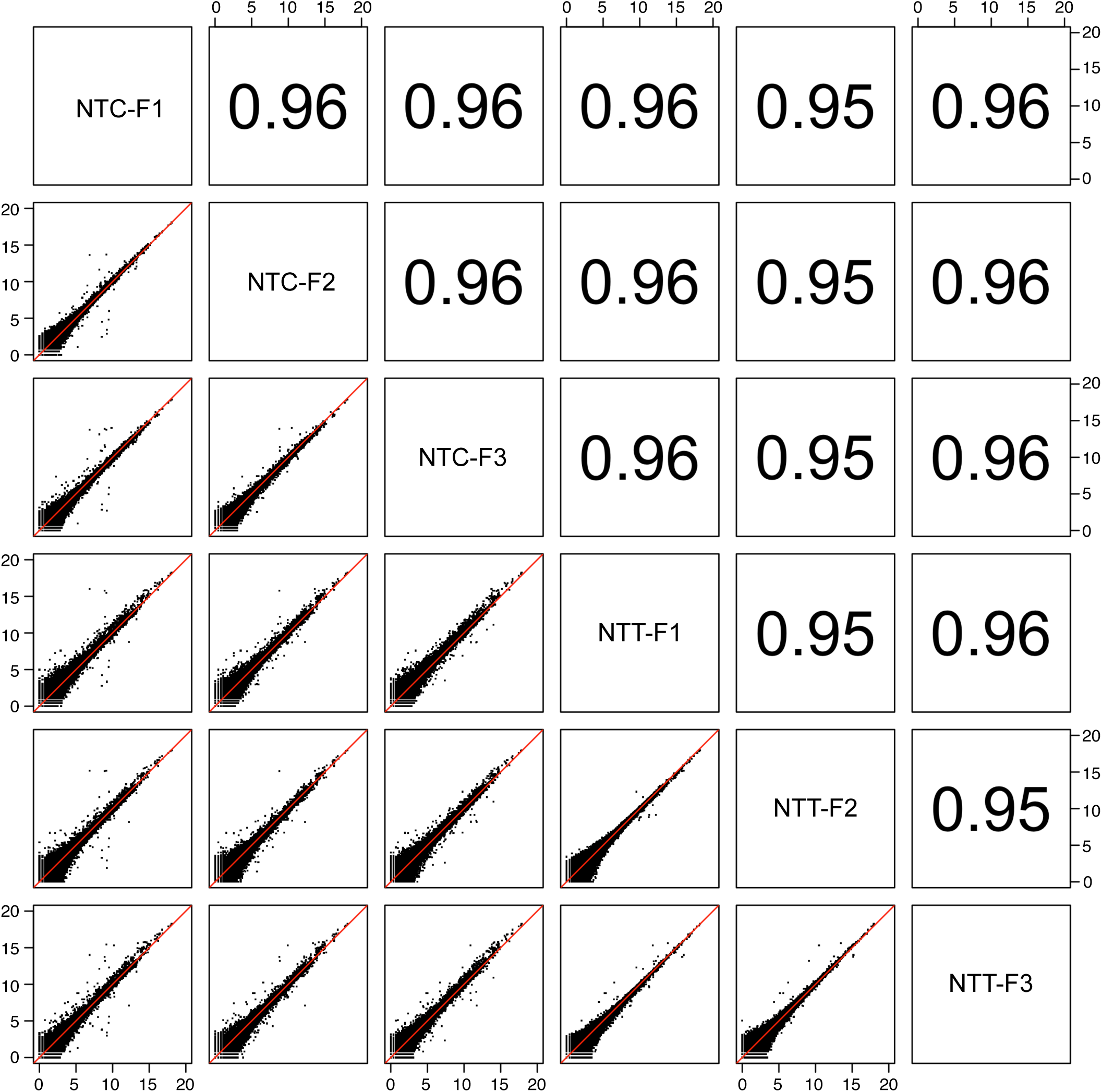
Scatter plot matrix of technical replicates in TGIRT-seq datasets of fragmented UHR RNAs plus ERCC spike-ins using either the NTC or NTT adapters. The x- and y-axes show DESeq2 normalized counts (log_2_ scale). Spearman’s correlation coefficient (ρ) are indicated in the upper right boxes.

**FIGURE S2.**
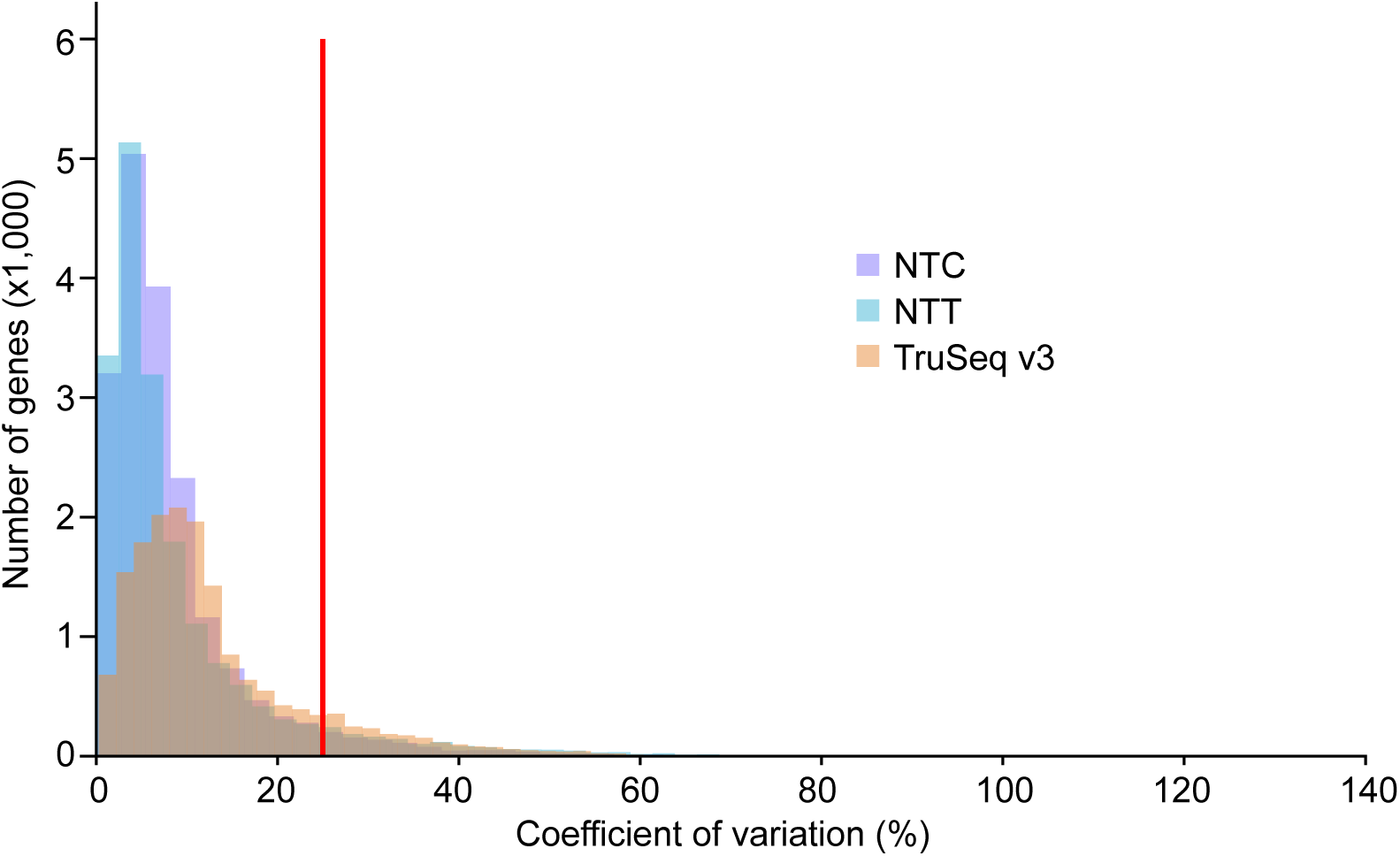
Histogram of coefficients of variation of protein-coding gene transcripts and ERCC spike-ins for TGIRT-seq of ribodepleted fragmented UHR RNA samples using the NTC or NTT adapters compared to those for TruSeq v3 (Li et al. 2014; SRA accession number SRP026126). Coefficients of variation were computed for each gene among technical replicates (n = 3 for NTC and NTT; n = 4 for TruSeq v3) and plotted as a histogram (bin size = 2.5). Only genes with DESeq2 (Love and Huber 2014) mean normalized counts >10 were included (n = 18,457, 18,459, and 16,723 for NTC, NTT, and TruSeq v3, respectively).

**FIGURE S3.**
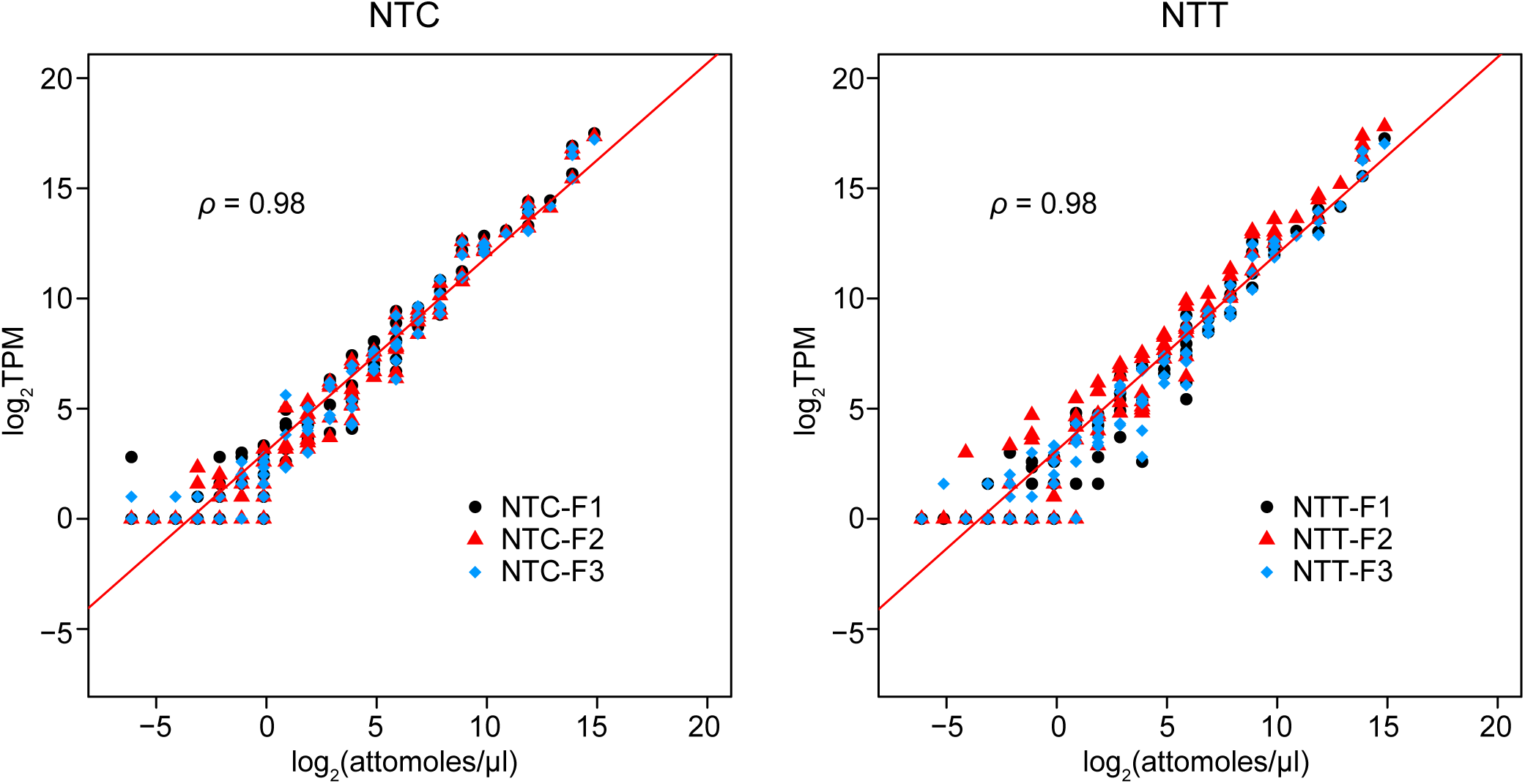
Scatter plot of ERCC spike-ins from fragmented UHR RNAs using either the NTC (left) or NTT (right) adapters. The x- and y-axes show the normalized counts (TPM, log_2_ scale) and expected concentration (attomoles/μl), respectively, for each ERCC spike-in. Spike-ins from the three technical replicates are shown in different colors and symbols. Spearman’s correlation coefficients (ρ) are indicated at the upper left.

**FIGURE S4.**
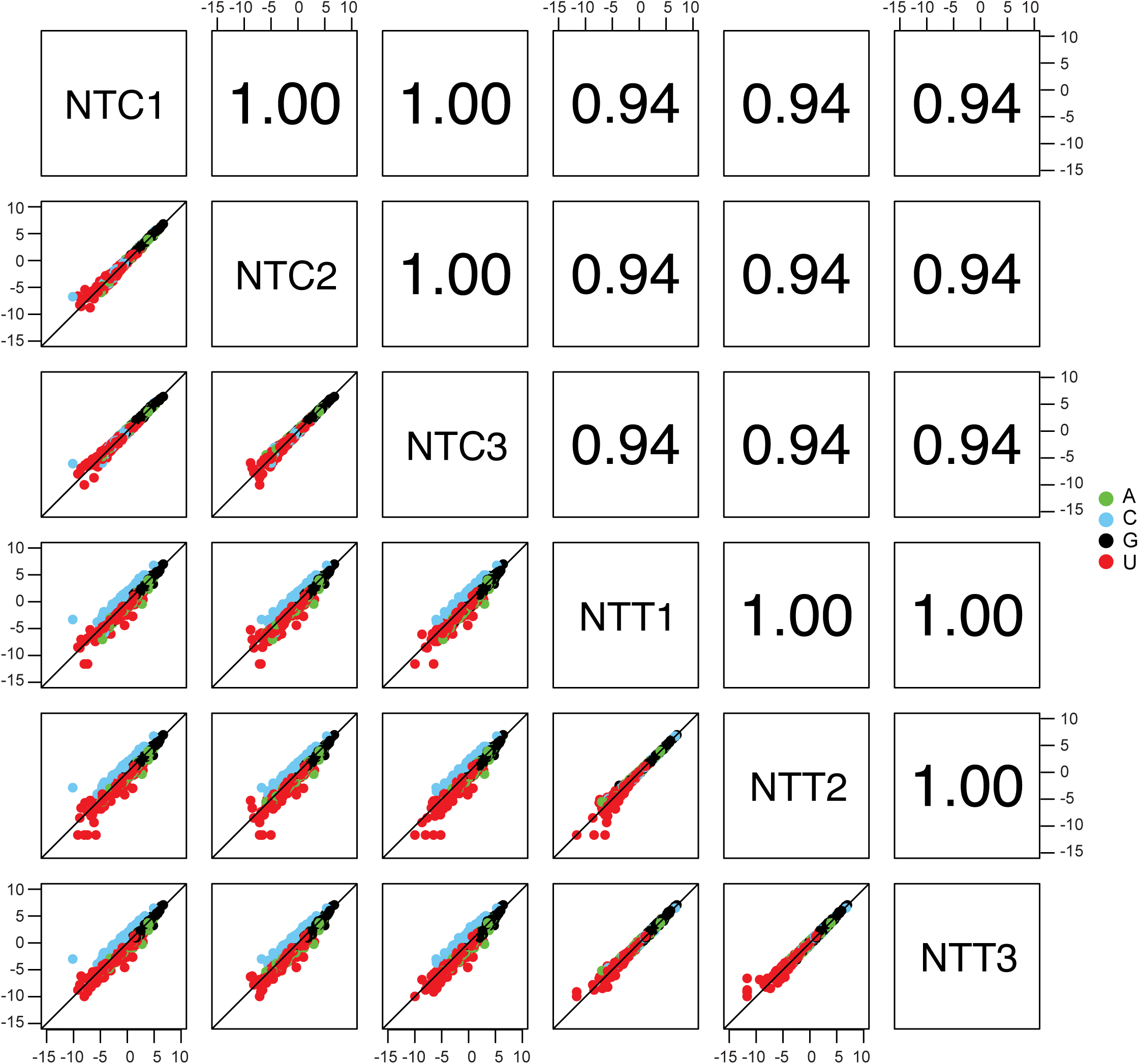
Scatter plot matrix comparing TGIRT-seq datasets obtained from the Miltenyi miRXplore miRNA reference set using either the NTC or NTT adapters (datasets NTC1-3 and NTT1-3, respectively). The x- and y-axes show median normalized counts (log_2_ scale). Spearman’s correlation coefficients (ρ) are indicated in the upper right boxes. miRNAs with different 3’ end nucleotides are colored coded (A, green; C, blue; G, black; U, red).

**FIGURE S5.**
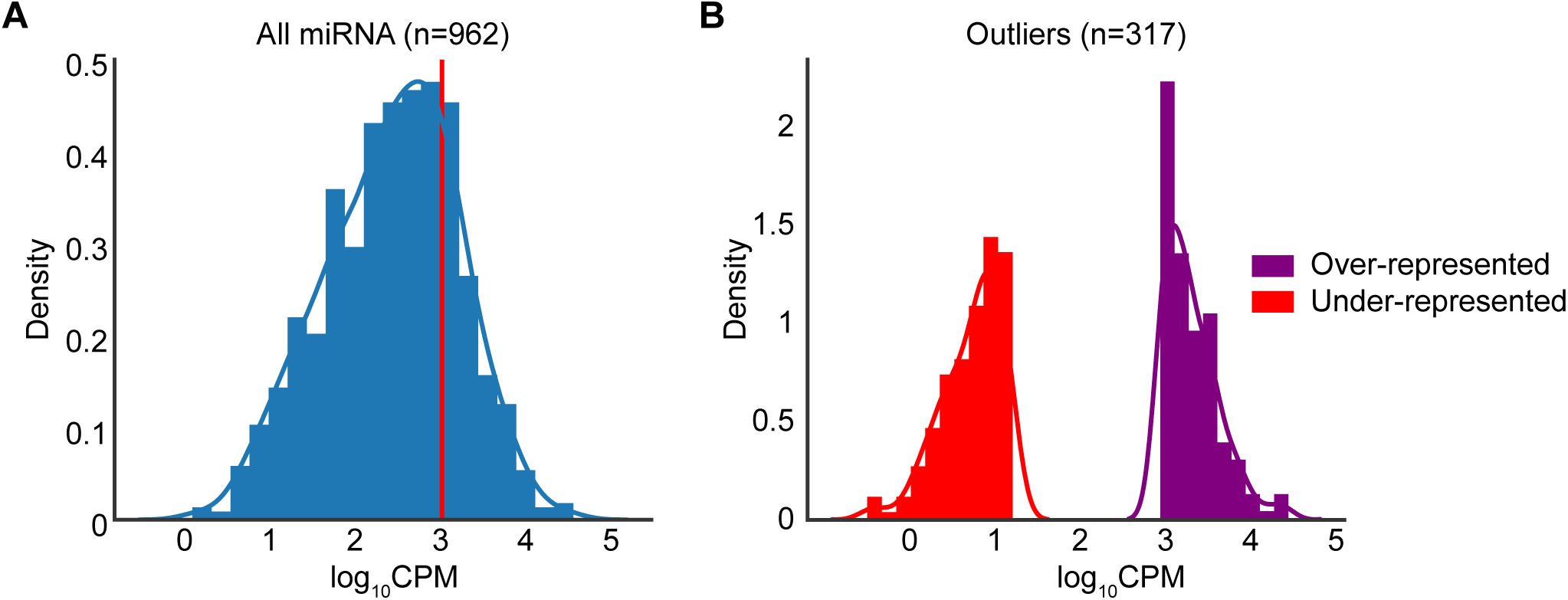
miRNA grouped by representation of sequences TGIRT-seq datasets obtained from the Miltenyi miRXplore miRNA reference set with the NTT adapter. (*A*) Histogram showing all 962 reference set miRNAs as a function of their log_10_CPM in combined TGIRT-seq datasets for the three technical replicates obtained using the NTT adapter. The red line indicates expected CPM value for the equimolar mix of 962 miRNAs. The histogram was computed using bin size of 0.45 in a log_10_CPM scale. (*B*) Histogram of under-represented (red) and over-represented (purple) miRNAs with log_10_CPM at least one standard deviation lower or higher than the mean log_10_CPM for all miRNAs in the reference set. The histogram was computed using bin size of 0.15 in a log_10_CPM scale.

**FIGURE S6.**
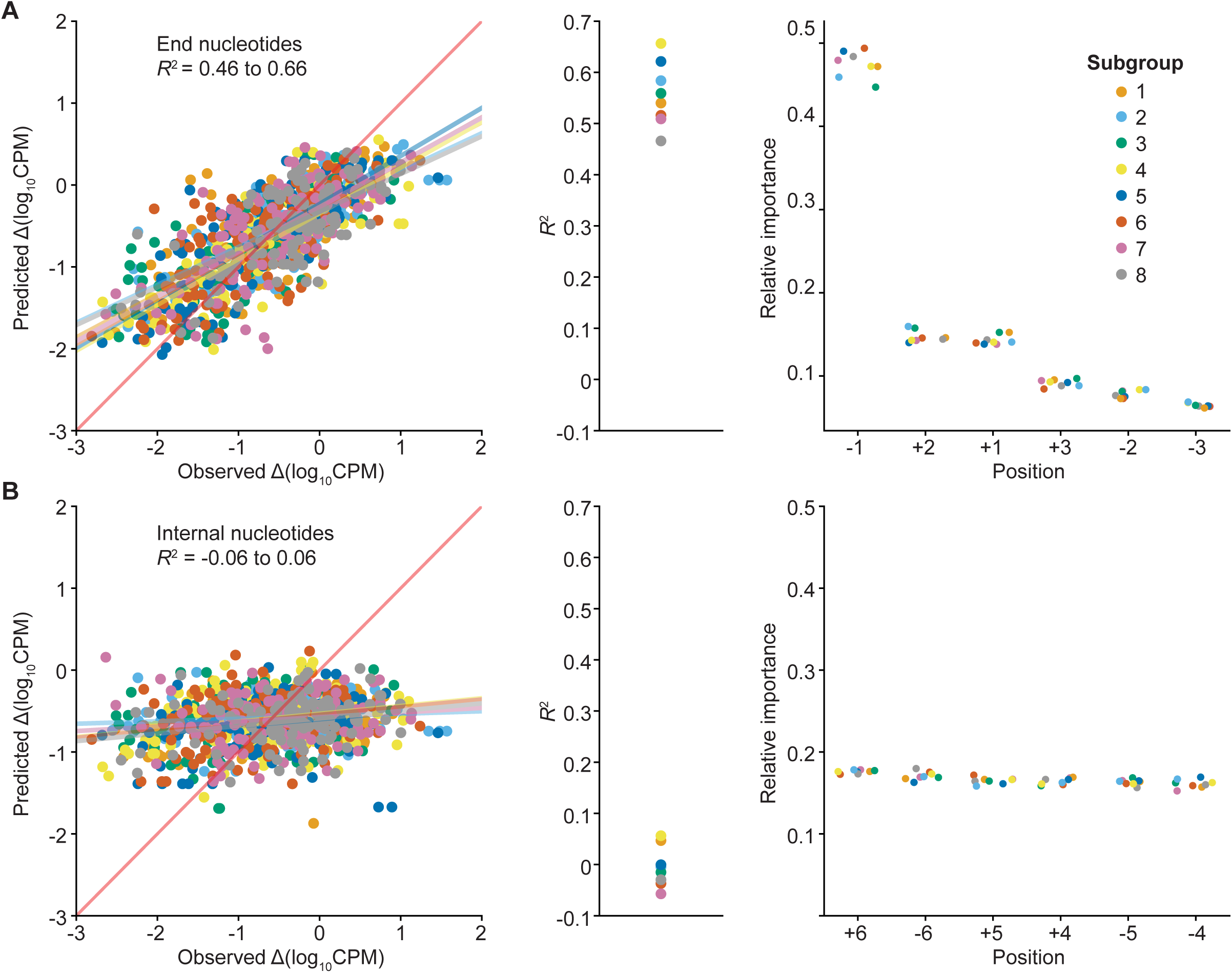
*k*-fold cross-validations of random forest regression bias-correction models based on the 5’- and 3’-end or internal nucleotide positions of miRNAs in combined TGIRT-seq datasets obtained with the NTT adapter for the Miltenyi miRXplore miRNA reference set. (*A*) shows results for models trained on the first 3 nucleotide positions from the 5’ and 3’ end of each miRNA (positions +1 to +3 and −1 to −3, respectively), and (*B*) shows results for models trained on the next 3 internal positions (positions +4 to +6 and −4 to −6, respectively). In each case, the 962 miRNAs in the dataset were randomly partitioned into 8 subgroups (120 or 121 miRNAs per group), and random forest regression models were trained against the observed measurement errors for each miRNA in a dataset comprised of 7 of the subgroups (Δlog_10_CPM: the difference between the observed and expected log_10_CPM for that miRNA) and tested on the remaining subgroup. The plots at the left show the predicted measurement errors (Δlog_10_CPM predicted by the random forest regression model) plotted against the observed measurement errors (Δlog_10_CPM obtained directly from sequencing data) for each miRNA color-coded by the subgroup on which the model was tested. The fitted linear regressions for each model were plotted as similarly color-coded solid lines, with the red diagonal line indicating hypothetical perfect prediction with slope = 1 and y-intercept = 0. The plots in the middle show *R*^2^ values for each of the models, and the plots at the right show the relative importance of each nucleotide position in each model, in each case color-coded by the subgroup on which the model was tested.

**FIGURE S7.**
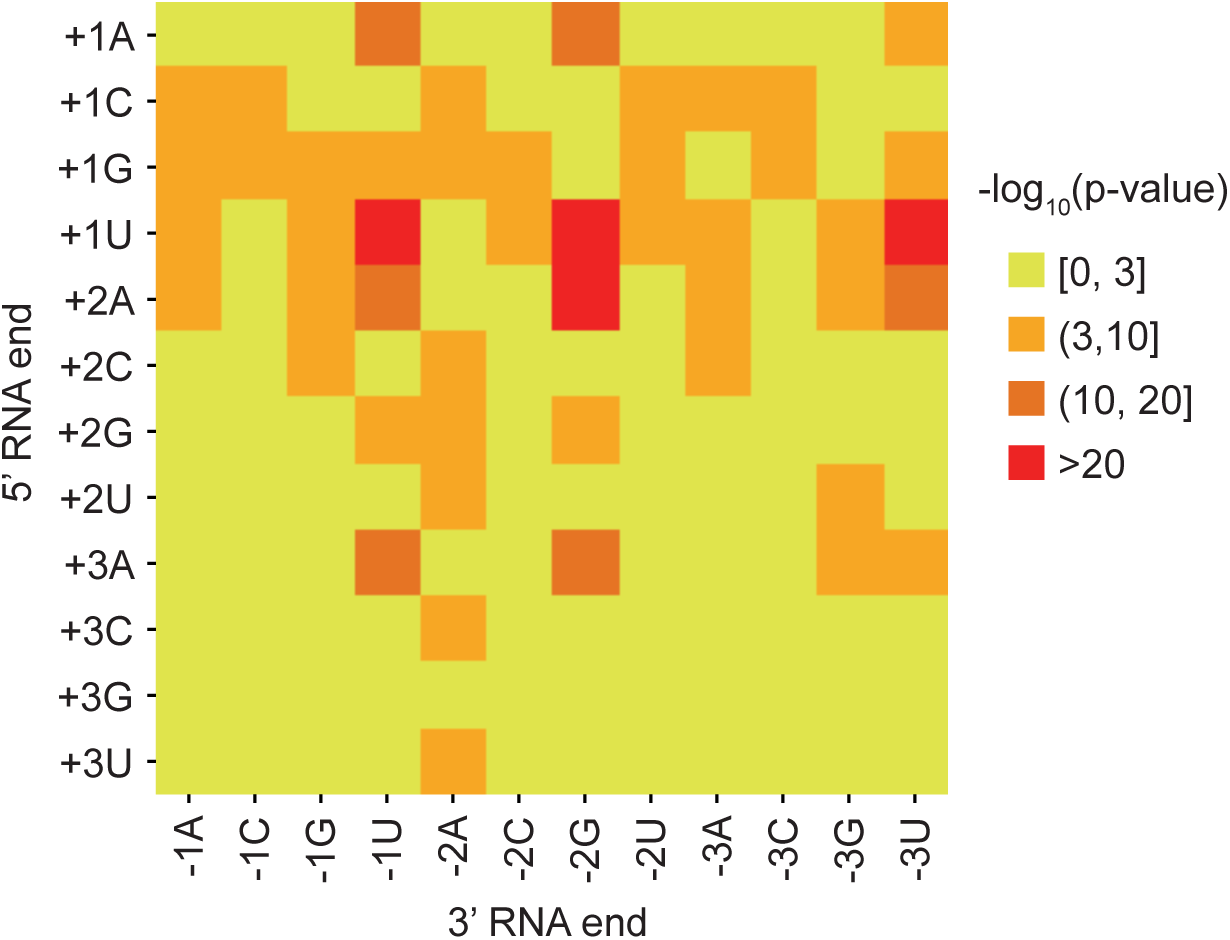
Sequence correlations between 5’- and 3’-end nucleotide positions in reference set miRNAs. Co-occurrences of nucleotide pairs from 5’ and 3’ ends (position N+1 to +3 and N-1 to −3, respectively) of the miRNAs in the Miltenyi miRXplore reference set were counted, and each pair was tested against a uniform distribution (16 different nucleotide patterns per position pair) using a χ^2^-test. Minus log_10_ p-values were adjusted for multiple comparisons by using the method of (Benjamini and Hochberg 1995) from the χ^2^-test and plotted as heat map for each of the paired nucleotide position color-coded by significance as indicated in the color bar.

**Table S1.**
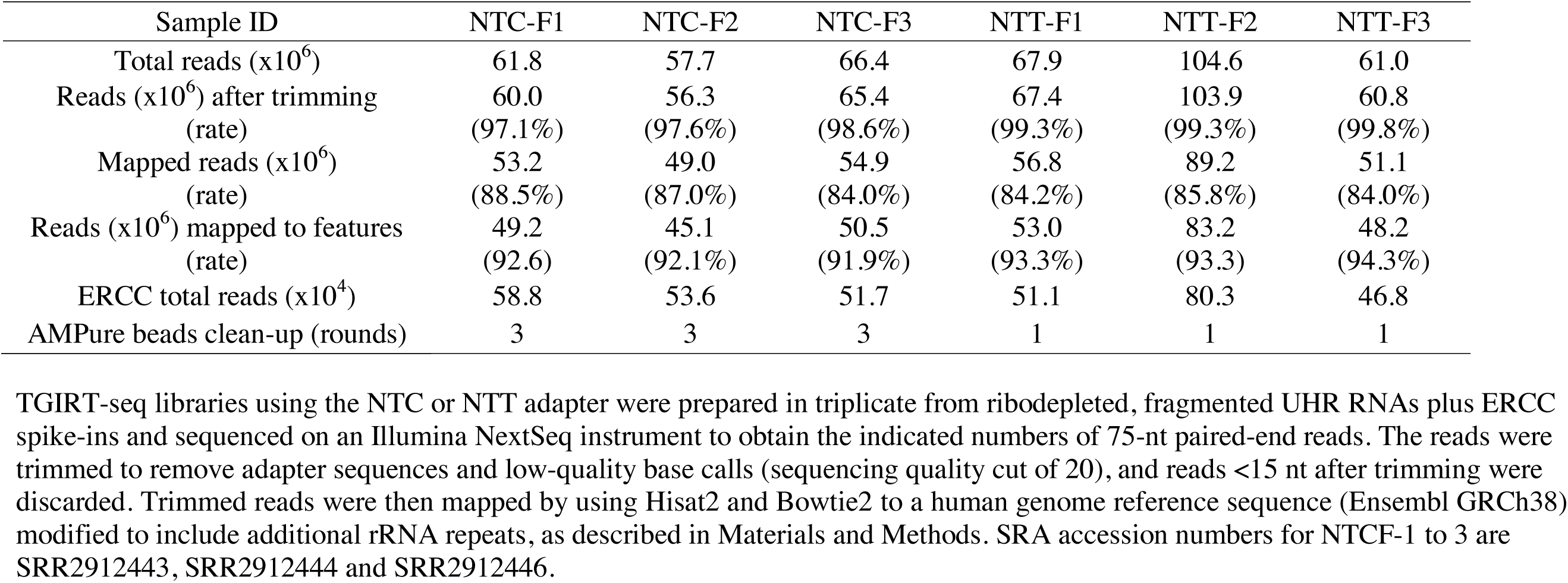
Read statistics and mapping for TGIRT-seq of the ribodepleted, fragmented UHR RNAs.

**Table S2.** Read statistics and mapping for TGIRT-seq of the Miltenyi miRXplore miRNA reference set.

| Sample ID | NTC1 | NTC2 | NTC3 | NTT1 | NTT2 | NTT3 |
| --- | --- | --- | --- | --- | --- | --- |
| Total reads ($\times 10^6$) | 14.1 | 12.3 | 13.3 | 15.0 | 15.6 | 18.2 |
| Reads ($\times 10^6$) after trimming<br>(rate) | 11.6<br>(82.2%) | 10.1<br>(81.9%) | 11.6<br>(87.6%) | 12.8<br>(85.2%) | 13.4<br>(85.9%) | 16.1<br>(88.1%) |
| Mapped reads ($\times 10^6$)<br>(rate) | 10.1<br>(87.1%) | 8.9<br>(88.1%) | 10.2<br>(87.7%) | 11.9<br>(93.1%) | 12.4<br>(93.0%) | 15.0<br>(93.4%) |
| Uniquely mapped reads ($\times 10^6$)<br>(rate) | 7.5<br>(74.1%) | 6.1<br>(68.8%) | 6.4<br>(62.8%) | 10.4<br>(87.9%) | 10.7<br>(85.9%) | 13.0<br>(86.6%) |
| AMPure beads clean-up (rounds) | 4 | 4 | 4 | 1 | 1 | 1 |

TGIRT-seq libraries using either the NTC or NTT adapter were prepared in triplicate from the Miltenyi miRXPlore miRNA reference set and sequenced on an Illumina NextSeq instrument to obtain the indicated numbers of 75-nt paired-end reads. The reads were trimmed to remove adapter sequences and low-quality base calls (sequencing quality cut of 20), and reads <15 nt after trimming were discarded. Trimmed reads were then mapped by using Bowtie2 to a 962 miRNA reference sequences, as described in Materials and Methods.

